# The Vps13-like protein BLTP2 is pro-survival and regulates phosphatidylethanolamine levels in the plasma membrane to maintain its fluidity and function

**DOI:** 10.1101/2024.02.04.578804

**Authors:** Subhrajit Banerjee, Stephan Daetwyler, Xiaofei Bai, Morgane Michaud, Juliette Jouhet, Shruthi Madhugiri, Emma Johnson, Chao-Wen Wang, Reto Fiolka, Alexandre Toulmay, William A. Prinz

**Affiliations:** Department of Cell Biology, University of Texas Southwestern Medical Center, Dallas, TX, USA; Lyda Hill Department of Bioinformatics, University of Texas Southwestern Medical Center, Dallas, TX, USA; Department of Biology, University of Florida, Gainesville, FL, USA; Genetics Institute, University of Florida, Gainesville, FL, USA; Université Grenoble Alpes, CNRS, CEA, INRAE, IRIG, LPCV, Grenoble, France; Department of Life Sciences, National Cheng Kung University, Tainan, Taiwan

## Abstract

Lipid transport proteins (LTPs) facilitate nonvesicular lipid exchange between cellular compartments and have critical roles in lipid homeostasis^1^. A new family of bridge-like LTPs (BLTPs) is thought to form lipid-transporting conduits between organelles^2^. One, BLTP2, is conserved across species but its function is not known. Here, we show that BLTP2 and its homolog directly regulate plasma membrane (PM) fluidity by increasing the phosphatidylethanolamine (PE) level in the PM. BLTP2 localizes to endoplasmic reticulum (ER)-PM contact sites^3^ ^4, 5^, suggesting it transports PE from the ER to the PM. We find BLTP2 works in parallel with another pathway that regulates intracellular PE distribution and PM fluidity^6, 7^. BLTP2 expression correlates with breast cancer aggressiveness^8–10^. We found BLTP2 facilitates growth of a human cancer cell line and sustains its aggressiveness in an in vivo model of metastasis, suggesting maintenance of PM fluidity by BLTP2 may be critical for tumorigenesis in humans.

## Results and Discussion

Research in plants and *Drosophila* revealed that mutations in BLTP2 reduce growth and cause defects in vesicular trafficking and phosphoinositide metabolism^2^. A recent study suggests BLTP2 operates at organelle contact sites and inhibits ciliogenesis in retinal pigment epithelial-1 cells ^11^. The finding that mice lacking BLTP2 are embryonic lethal suggests BLTP2 has an essential role in mammalian development^12^ (https://www.mousephenotype.org/data/genes/MGI:1919753). To investigate BLTP2’s importance in cancer, we examined data from the Cancer Dependency Map (DepMap) and found that *BLTP2* is highly expressed and near essential in invasive breast cancer cell lines (Fig. 1a), together suggesting a critical role in breast cancer progression^13^ ^14^. We found that CRISPR/Cas9 mediated knock-out of *BLTP2* (BLTP2-KO) in the cervical cancer cell line HeLa also reduced its rate of proliferation (Fig. 1b). Together, these results suggest BLTP2 is important for cell growth in mammals and may be particularly important to sustain rapid proliferation of some cancer cells.

**Figure 1.**
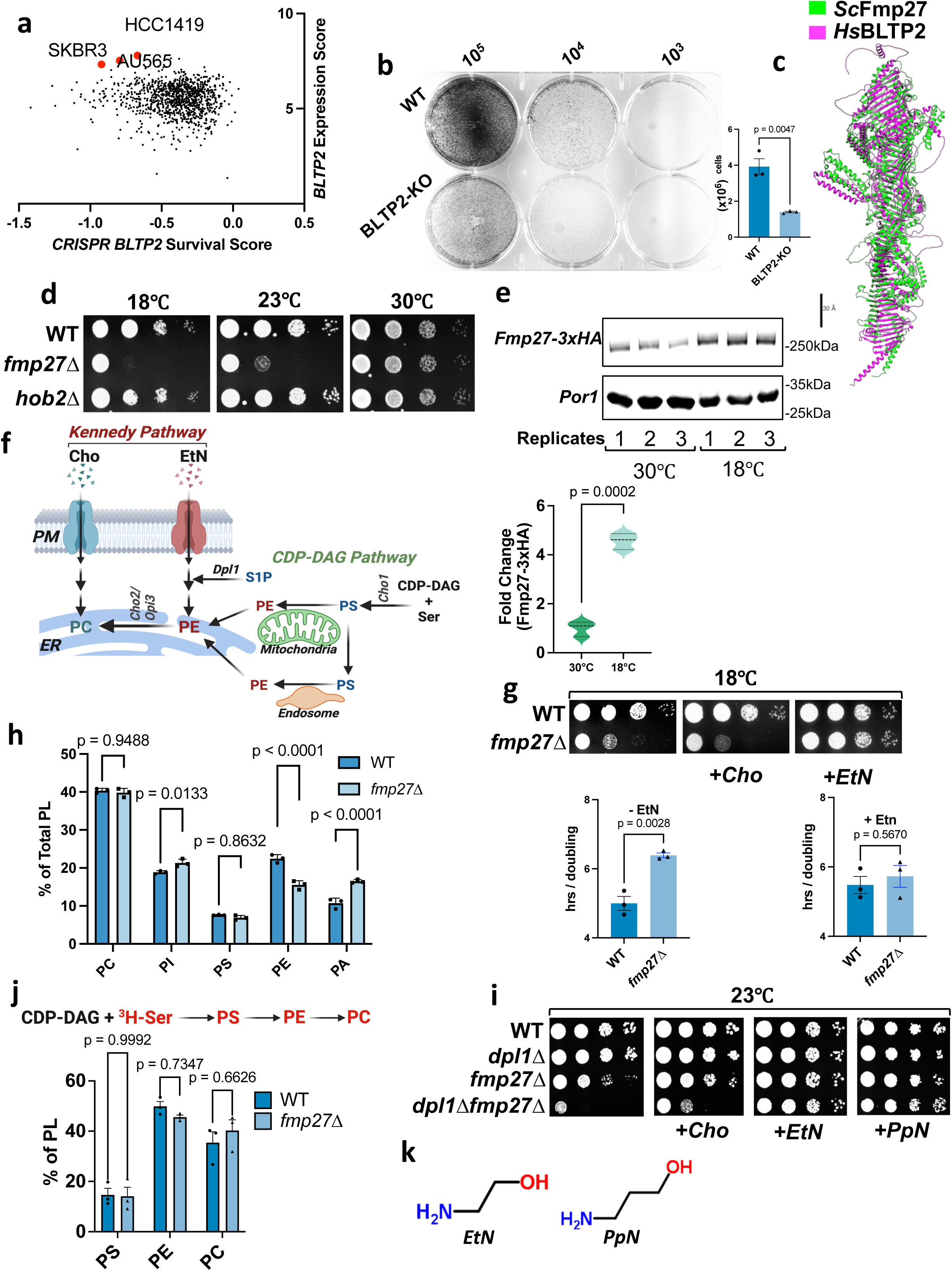
PE rescues the cold sensitivity of yeast lacking the BLTP2 homolog, Fmp27. a. BLTP2 mRNA expression score (between 0.5 – 10) plotted against a survival score (–1 to +1) of about 1000 cancer cell lines with a deletion of BLTP2. Data were collected from DepMap and plotted manually. b. Crystal violet-stained colony forming units of HeLa cells with the indicated genotypes plated at the cell number indicated on the top (left). Growth rate of WT and BLTP2-KO cells (n=3) (right). Mean cell number±SEM is plotted in a histogram; p-value from an unpaired t-test. c. The alphafold predicted secondary structure of yeast Fmp27 (green) and human BLTP2 (magenta) was compared and aligned using the Matchmaker plug-in of UCSF Chimera. Scale bar = 30Å. d. Serial dilutions of yeast strains were spotted on SC medium without EtN supplementation. Plates were grown for 3 days. e. Western blot (top) of genomically expressed Fmp27-3xHA and Por1 (control) from n=3 samples (indicated by replicate no.). Samples are from yeast grown at the indicated temperatures. Quantification of the Fmp27 protein at each temperature normalized to Por1 protein amounts is depicted in the box-violin plot with a p-value from an unpaired t-test (bottom). f. Schematic for the Kennedy and CDP-DAG pathways of PE and PC biosynthesis drawn using biorender.com. g. Serial dilutions yeast strains spotted on SC medium without, or with 4mM choline (Cho) or 4mM ethanolamine (EtN) supplementation (top). The plates were grown at 18°C for 3 days. Doubling time (hrs/doubling) of the indicated strains grown at 18°C, with or without EtN supplementation, in a 96 well plate; histograms show mean± SEM (n=3) (bottom); p-value from unpaired t-test. h. The relative amount of the five major phospholipids (PL) in cells grown at 18°C and labeled to steady-state with [^3^H]palmitate (PS, phosphatidylserine; PI, phosphatidylinositol; PA, phosphatidic acid); mean± SEM (n=3), p-values from unpaired t-test. i. Serial dilution of yeast strains spotted on SC medium without or with Cho, EtN, or PpN and incubated at 23°C for 3 days. j. Scheme of PS, PE, and PC production by exogenous [^3^H]serine (top). Bar graph shows the relative abundance of radiolabeled PLs after growth for 1 hr at 18°C (bottom); mean± SEM (n=3); p-values from unpaired t-test. k. Chemical structures of EtN and propanolamine (PpN).

To get more insight into BLTP2 function, we turned to *S. cerevisiae* (hereafter yeast), which has two BLTP2 homologs: Fmp27 and Hob2^3^. Human BLTP2 and Fmp27 have similar predicted structures (Fig. 1c). Since the functions of Fmp27 and Hob2 are unknown, we sought to identify phenotypes for cells lacking either protein (*fmp27Δ* and *hob2Δ*, respectively) and observed *fmp27Δ* cells grew poorly at low temperatures (Fig. 1d). Consistent with a role for Fmp27 in cold adaptation, levels of endogenous Fmp27 increased four-fold when cells are grown at low temperature (18°C; Fig. 1e). We ruled out that growth at low temperature alters the localization of Fmp27; it localizes to ER-PM contact sites at both 30°C^4^ and 18°C (Fig. S1a).

While cells lacking Fmp27 grew poorly at low temperatures on Synthetic Complete (SC) media (Fig. 1d), growth was normal on Yeast Peptone Dextrose (YPD; Fig. S1b). This suggested SC lacks nutrients present in YPD that support the growth of *fmp27Δ* cells at low temperatures. Two nutrients present in YPD but absent from SC are ethanolamine (EtN) and choline (Cho). Exogenous EtN and Cho support the biosynthesis of phosphatidylethanolamine (PE) and phosphatidylcholine (PC) by the Kennedy pathway (Fig. 1f). Yeast does not synthesize Cho and the only source of endogenous EtN is EtN-phosphate made by degrading dihydrosphingosine phosphate, catalyzed by Dpl1^15^. When exogenous Cho and EtN are not present, PC and PE are synthesized from cytidine diphosphate diacylglycerol (CDP-DAG; Fig 1f). We found that the growth defect of cells lacking Fmp27 in SC at 18°C is corrected by supplementation of EtN but not Cho to the medium (Fig. 1g), suggesting *fmp27D* cells require EtN to grow at low temperatures; growth at 30°C was not affected by exogenous EtN (Fig. S1c).

Since yeast only uses EtN to produce PE, *fmp27Δ* cells may grow poorly at low temperatures because they have insufficient PE. Consistent with this inference, these cells have reduced PE levels when grown at 18°C (Fig 1h). To obtain further evidence that *fmp27Δ* cells have insufficient PE, we used strains lacking Dpl1, which produce low levels of PE in SC media^16^ ^15^. Cells lacking both Dpl1 and Fmp27 proliferated more poorly at 23°C than cells lacking only Fmp27, and growth was restored by exogenous EtN but not Cho (Fig. 1i), suggesting the growth defect of *fmp27Δ* cells is caused by altered PE homeostasis.

We ruled out that cells lacking Fmp27 have defects in producing PE via the CDP-DAG pathway (Fig. 1f). In this pathway, cells produce phosphatidylserine (PS) that is decarboxylated to form PE, which is methylated to generate PC. To estimate the rate of flux through the CDP-DAG pathway in vivo, we labeled cells with [^3^H]serine at 18°C for 60 minutes and determined the amount of radiolabel in PS, PE, and PC. We found no significant difference (Figs. 1j, S1d). We also ruled out the possibility that cells lacking Fmp27 have defects in producing PS, which would be expected to decrease rates of PE production via the CDP-DAG pathway. If this were correct, the growth defect of cells lacking Fmp27 would be corrected by overproducing PS synthase (Cho1), but this was not observed (Fig. S1e).

Since cells lacking Fmp27 do not seem to have defects producing PE by the CDP-DAG pathway or using PE to produce PC, we wondered whether they have defects in other metabolic processes requiring PE. One is the production of glycosylphosphatidylinositol (GPI) anchors, which requires three molecules of PE per GPI^17^. Cells with defects in GPI-anchor biosynthesis are hypersensitive to calcofluor white^18^. However, we found that cells lacking Fmp27 are not hypersensitive to CFW (Fig. S1f), suggesting they do not have a defect in GPI-anchor biosynthesis. This conclusion is further supported by our analysis of whether *fmp27Δ* cells are hypersensitive to tunicamycin, which induces ER stress. We previously showed that yeast cells lacking the Vps13 family protein Csf1 have defects in GPI anchor biogenesis and are hypersensitive to tunicamycin^4^, but cells lacking Fmp27 do not (Fig. S1g). We conclude that cells lacking Fmp27 do not have defects in PE-requiring metabolism.

We next considered whether the biophysical properties of PE are important for sustaining the growth of cells lacking Fmp27 at low temperatures. PE, unlike the other abundant phospholipids in eukaryotic cells, has a high negative monolayer spontaneous curvature and forms hexagonal phase structures rather than membrane bilayers, properties thought to be necessary to sustain membrane dynamics and fluidity^19^. A previous study showed that exogenous propanolamine (PpN; Fig. 1k) is used in place of EtN by yeast to produce phosphatidylpropanolamine, a phospholipid with the biophysical properties of PE but that cannot be converted to PC^16^. We found that exogenous PpN improves the growth of *fmp27D* cells at 23°C (Fig. 1i), indicating that the biophysical properties of PE are critical for the growth of these cells.

These properties are important for the role of PE in maintaining membrane fluidity, which is determined, in part, by the ratio of PE to bilayer-forming lipids such as PC (the most abundant lipid in most cellular membranes^20^). Cells also maintain membrane fluidity by modulating the saturation of acyl chains in membrane lipids and other mechanisms, a process known as homeoviscous adaptation^21^. Because a role for another BLTP protein in homeoviscous adaptation has been suspected ^22, 23^, we wondered whether Fmp27 might regulate membrane fluidity at low temperatures by modulating the ratio of PE to PC. We first determined the PE:PC ratio for wild-type (WT) and *fmp27Δ* cells grown at 30°C and 18°C. When wild-type cells are grown in a medium without EtN, the PE:PC ratio increases significantly at 18°C (Fig. 2a), which is likely a mechanism of homeoviscous adaptation. In contrast, the PE:PC ratio of cells lacking Fmp27 does not change when they are grown at 18°C. The ratio is also significantly lower than wild-type at 18°C (Fig. 2a, left). This suggests that *fmp27Δ* cells fail to alter PE:PC ratio at low temperatures when cells are grown without EtN. When EtN was added to the medium, there was no significant difference in the ratio of PE:PC in wild-type and *fmp27Δ* cells (Fig. 2a, right).

**Figure 2.**
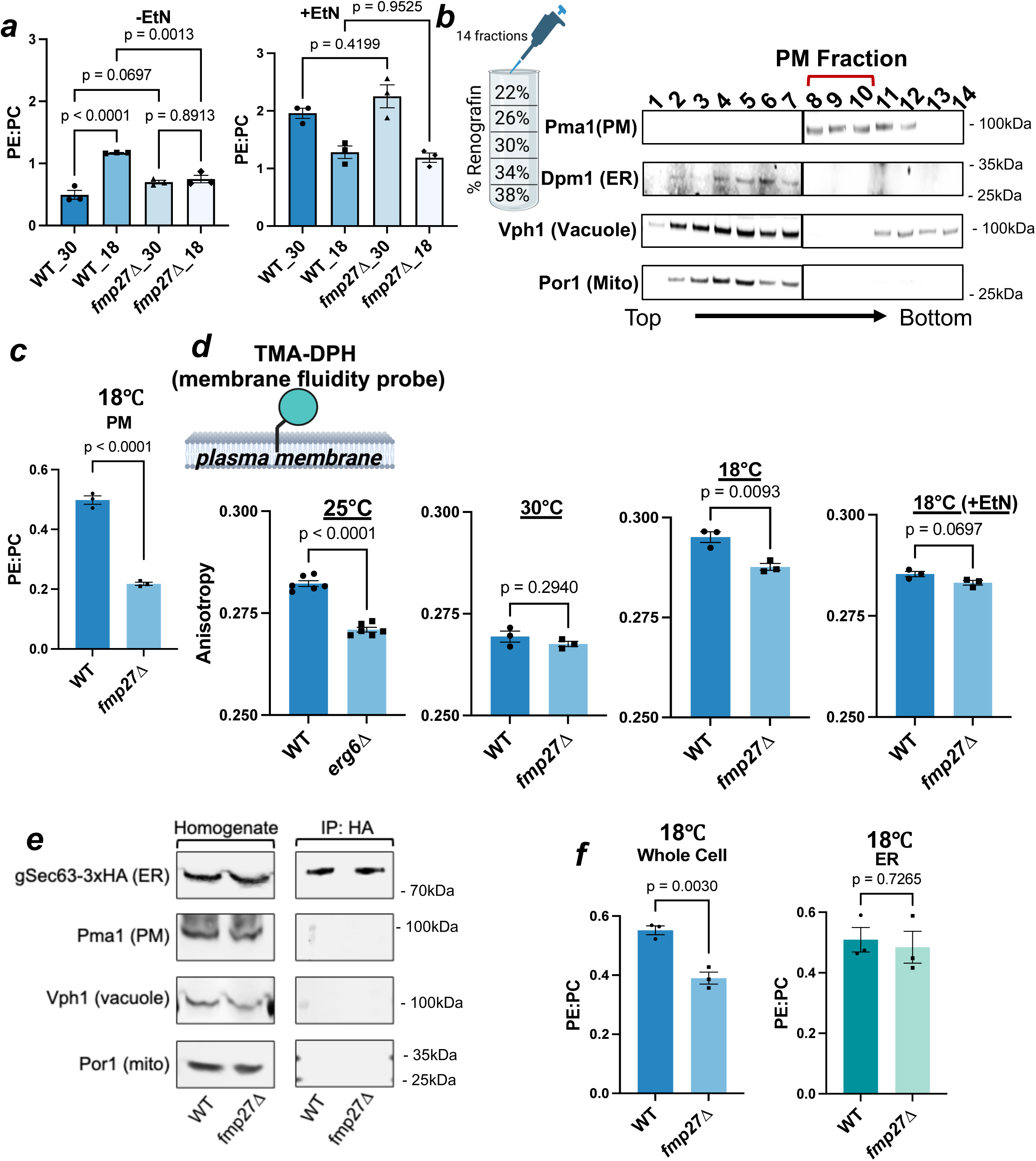
Cells lacking Fmp27 have reduced PE in the PM and increased PM fluidity at low temperature. a. PE:PC ratios from cells grown at the indicated temperatures with or without 4mM EtN; mean± SEM (n=3); p-values from one-way ANOVA. PLs measured using ELSD. b. Scheme showing how Renografin-76 density gradients were used to obtain PM-enriched fractions of cell lysates (left). Western blot of indicated proteins from equal volumes of each of the 14 fractions separated by SDS-PAGE; numbers indicate fraction, PM fractions indicated with red bracket. Images from different blots separated by borders. c. PE:PC ratios from PM-enriched fraction from strains grown at 18°C; mean± SEM (n=3) of PE:PC; p-values from unpaired t-test. PLs measured using ELSD. d. Cartoon of the fluidity probe TMA-DPH (circle) in the outer leaflet of the PM (top; generated using biorender.com). TMA-DPH anisotropy from yeast strains grown at indicated temperatures with or without EtN (bottom); mean± SEM (n≥ 3); p-values from unpaired t-tests. e. Western blots of immunopurified ER from yeast strains expressing Sec63-3xHA and grown at 18°C. Blots of whole cell homogenates (left) and after immunopurification of membranes using an anti-HA antibody (IP:HA; right). f. PE:PC ratios of whole cell lysates and ER membranes as in e; mean± SEM (n= 3); p-values from unpaired t-tests. Cells were labeled to steady-state with [^3^H]palmitate to allow measurement of PLs.

Since Fmp27 localizes to ER-PM contact sites, we determined whether it is particularly important for maintaining the ratio of PE:PC in the PM. Using density gradient centrifugation to isolate PM-enriched fractions^24^ from cells grown at 18°C (Fig. 2b), we found that the PE:PC ratio of the PM was ∼2.5 fold higher in wild-type cells than in cells lacking Fmp27 (Fig. 2c). This suggests Fmp27 regulates PE levels in the PM to regulate fluidity. To directly assess PM fluidity, we used TMA-DPH, which has been used to assess fluidity in the outer leaflet of the PM^25^. Previous studies have shown that yeast cells lacking Erg6 (*erg6Δ*), which have zymosterol rather than ergosterol as their primary sterol, have increased PM fluidity^25^ ^26^, a finding we confirmed (Fig. 2d). We next found that the PM fluidity of *fmp27Δ* cells is indistinguishable from wild-type when cells are grown at 30°C (Fig. 2d). In contrast, when grown at 18°C the PM fluidity of *fmp27Δ* cells was significantly higher than that of wild-type. This difference is eliminated by adding EtN to the growth medium (Fig. 2d). Together, these findings indicate that cells lacking Fmp27 fail to alter the ratio of PE:PC when grown at a low temperature and have increased PM fluidity. We speculate that Fmp27 transports PE from the ER to the PM to maintain the PE:PC ratio of the PM at low temperatures.

We wondered whether Fmp27 might also play a role in determining the PE:PC ratio of the ER. We immunopurified ER-derived membranes^27^ (Fig. 2e) from wild-type and *fmp27Δ* cells to measure PE:PC ratio and found no significant difference (Fig. 2f). Note that the PE:PC ratio for whole cell extracts for these experiments was different than those shown in Fig. 2a. This is probably a result of the different methods used to quantify phospholipids: an evaporative light-scattering detector (ELSD) was used to generate the results in Fig. 2a,c, while cells labeled with [^3^H]palmitate were used to generate the results shown in Fig. 2f. However, with both methods we consistently observed a significant reduction in the PE:PC ratio in *fmp27Δ* cells grown at 18°C. Together, these results suggest that Fmp27 specifically regulates the PE:PC ratio of the PM but not the ER.

We wondered whether Fmp27 controls membrane fluidity not only by affecting PE:PC ratio but also by altering the saturation of phospholipid acyl chains, perhaps by regulating acyl chain remodeling. To address this, we determined the relative abundance of glycerophospholipids species in wild-type and *fmp27Δ* cells grown at 30°C and 18°C, both in a media without (Fig. 3a) and with EtN (Fig. 3b). No major differences were found. Calculation of an unsaturation index of glycerophospholipids revealed that while the index increases significantly when cells are shifted from 30°C to 18°C, there was no difference between wild-type and *fmp27Δ* cells (Fig. 3c). Interestingly, the unsaturation index was not affected by growth temperature when EtN was in the medium (Fig. 3d), suggesting changes in acyl chain composition do not play a significant role in cold adaptation when PE is abundant. These findings suggest Fmp27 does not regulate phospholipid acyl chain remodeling or production. This conclusion was also supported by a determination of whether the growth of *fmp27Δ* cells was affected by inositol, an important regulator of glycerolipid metabolism in yeast^28^ (Fig. 3e). We found that the relative growth rates of *fmp27Δ* and wild-type cells at 18°C were the same regardless of the presence of inositol in growth media (Fig. 3f).

**Figure 3.**
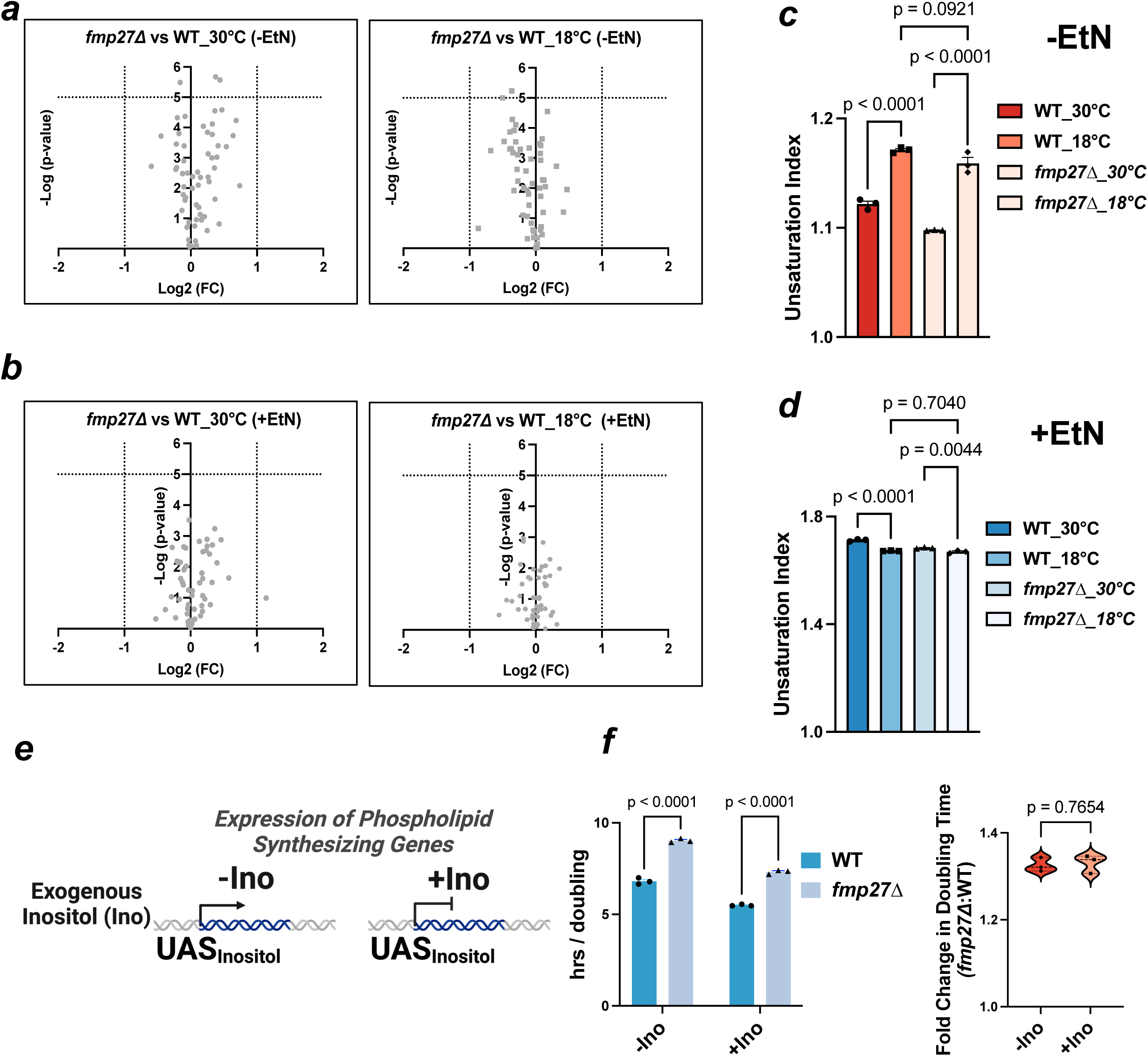
Acyl chain remodeling and lipid metabolism are not altered in cells lacking Fmp27. a,b. Volcano plots showing log fold changes (Log2 FC) of PL species against statistical significance (-Log p-value) in yeast strains grown in SC medium without (a) or with (b) EtN. c,d. Unsaturation indices PLs from yeast grown in SC without (c) or with (d) EtN; mean± SEM (n=3) values; p-values from one-way ANOVA. e. Scheme describing the role of inositol (Ino) in regulating transcription of PL biosynthetic genes in yeast. f. Doubling time of yeast strains grown at 18°C in SC lacking Ino or containing 100mM Ino. Results are shown as bar graphs (left) and as box-violin plots of relative difference between the strains (right; mean± SEM (n=3); p-values from unpaired t-tests.

We next determined whether BLTP2 has a similar function in the nematode model, *C. elegans*, which has one isoform of Fmp27. We named it *bltp-2*. We generated worms with a deletion of *bltp-2* and found they did not have growth or morphology defects (Fig. S2a-b). While a previous study showed that *Drosophila* lacking BLTP2 (called Hob) had reduced pupal size^29^, this was not found in *C. elegans bltp-2* mutant (Fig. S2a-c). However, we found that the body scale of *bltp-2* worms was significantly affected by RNAi-mediated knockdown of *pcyt-1,* which produces phosphate cytidyltransferase 1, the rate-limiting enzyme in the Kennedy pathway^30^ (Fig. S2c-e). This treatment also reduced brood size (Fig. S2f) and embryonic viability (Fig. S2g). Together, these findings suggest that in *C. elegans*, as in yeast, BLTP2 is important for cell growth and development when PE production by the Kennedy pathway is limited.

We found this is also true in human cells. The growth rate of HeLa BLTP2-KO cells in media with dialyzed FBS (dFBS), which does not contain EtN or Cho^31^, was improved by addition EtN but not Cho (Fig. 4a). Notably, addition of PpN also improved growth, suggesting that the biophysical properties of PE are important for the growth of BLTP2-KO cells when they do not have sufficient exogenous EtN to support robust PE production by the Kennedy pathway. Consistent with this, we measured PE and PC levels in cells grown in media with complete FBS (Fig. 4b) or dialyzed FBS (Fig. 4c) and found PE levels were significantly reduced in cells grown with dFBS. This was also found when we measured the ratio of PE to PC in the PM. We used an established method to obtain a PM-enriched fraction from attached cells^32^ (Fig. S3a), confirmed the fraction was PM-enriched (Fig. 4d), and found the PE:PC ratio was significantly reduced in PM from BLTP2-KO cells (Fig. 4e).

**Figure 4.**
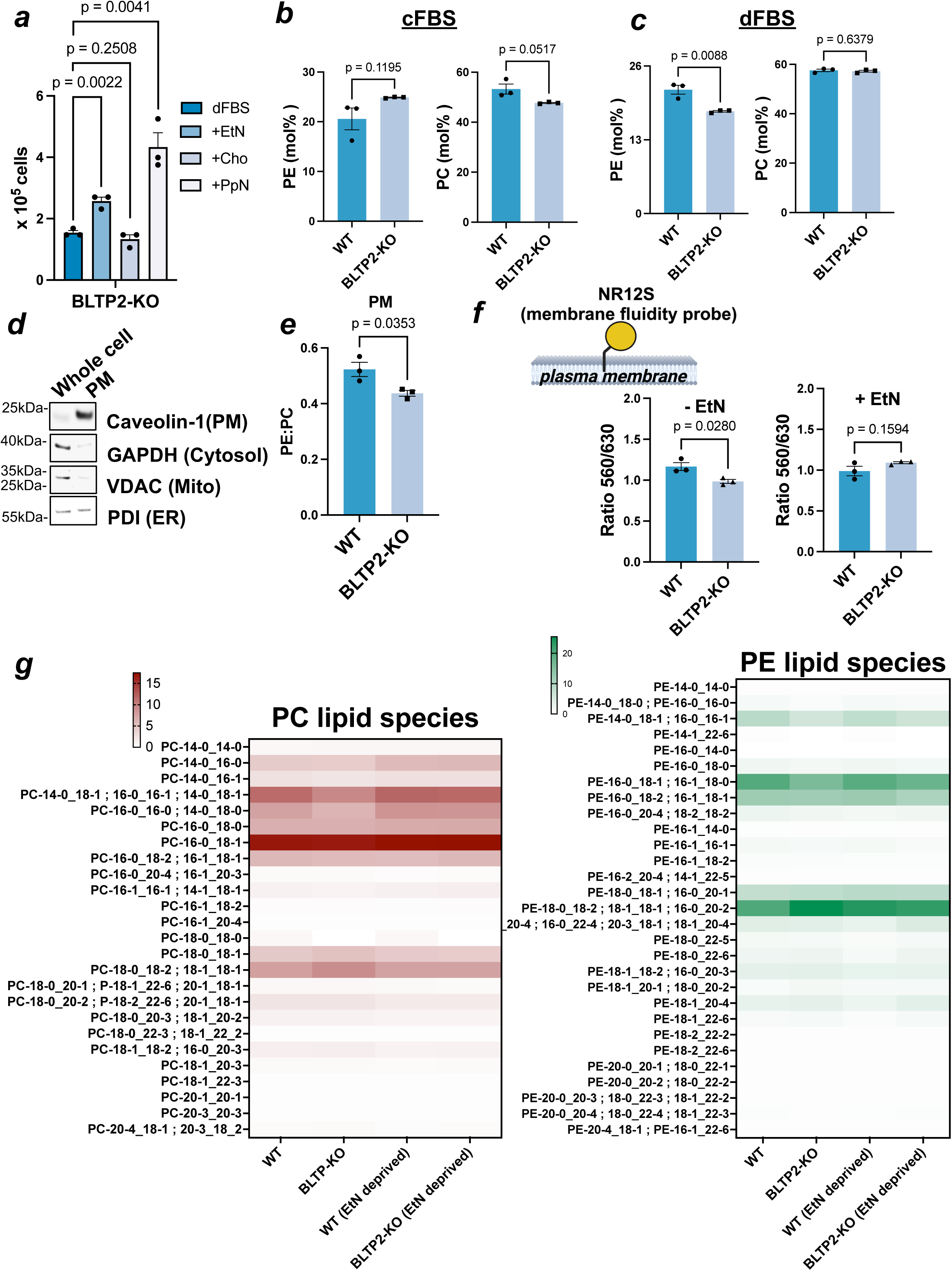
BLTP2-KO HeLa cells have PMs with a decreased PE:PC ratio and increased PM fluidity. a. Automated cell count assay for BLTP2-KO HeLa cells grown with DMEM containing dialyzed FBS without supplementation (dFBS) or with 10µM EtN, Cho, or PpN; mean± SEM (n=3) values; p-values are from unpaired t-tests. b-c Mol% of PLs from HeLa cells grown in DMEM with FBS (b) or dFBS (c) determined using LC-MS/MS; mean± SEM (n=3); p-values from unpaired t-tests. d. Western blots of whole cell lysates and PM-enriched fractions (PM) of HeLa cells grown in DMEM with dFBS. e. PE:PC ratios of PM-enriched fraction from HeLa cells in e; mean± SEM (n=3); p-values from unpaired t-tests. PL levels were determined by ELSD. f. Cartoon of the fluidity probe NR12S (circle) in the outer leaflet of the PM (top; generated using biorender.com). Ratio of NR12S fluorescence at 560nm and 630nm of HeLa strains grown in DMEM with dFBS without or with EtN supplementation (bottom); mean± SEM (n≥ 3); p-values from unpaired t-tests. g. Relative abundance of PE and PC species, determined by LE-MS/MS, from HeLa cells grown in DMEM with complete FBS or dFBS (EtN deprived) and shown as a heat map (0-lowest to 15-highest; mean of 3 independent replicates).

These findings prompted us to determine whether PM fluidity is reduced in BLTP2-KO cells as it is in yeast cells lacking Fmp27. We used the order sensitive dye NR12S, which has been used to measure fluidity in the outer leaflet of the PM^33^. We first confirmed we could measure membrane fluidity with this dye by showing that treating cells with methyl beta cyclodextrin (MbCD), which extracts cholesterol from cells, increases PM fluidity as previously shown^33^(Fig. S3b). We found that BLTP2-KO cells have increased PM fluidity when EtN is not in the medium (Fig. 4f). We speculated that human BLTP2, like its yeast homolog, controls PM fluidity primarily by increasing the ratio of PE:PC in the PM, but not by affecting phospholipid acyl chain saturation. Consistent with this, we found that the relative abundance of the major phospholipid species does not change in BLTP2-KO cells (Figs. 4g; S3e,f).

Together, our findings suggest that BLTP2 in humans, like its yeast homolog, regulates PM fluidity by increasing the amount of PE in the PM when PE synthesis by the Kennedy pathway is compromised by a lack of exogenous EtN. We speculate that BLTP2 transports PE from the ER to the PM. Consistent with this, we found that overexpressed BLTP2 localizes to ER-PM contact sites (Fig. S3e) like its homologs in yeast^4^ and *Drosophila*^3^.

We wondered whether BLTP2 in human cells operates together with a previously described pathway that regulates PM fluidity and PE metabolism. This pathway requires the PM protein Tram-Lag-CLN8 (TLC)-domain containing protein 1(TLCD1)^6^ ^7^. How TLCD1 functions is not well understood. We identified that TLCD1 is one of the top hits in a list of genes that are co-dependent with BLTP2^13^ ^14^ (Cancer Dependency Map Portal (RRID:SCR_017655); Fig. S4a). To confirm this, we treated HeLa cells in which TLCD1 has been knocked out (TLCD1-KO) with shRNAs targeting BLTP2 (shBLTP2) or a control, untargeted shRNA (shUT). We found that elimination of TLCD1 and BLTP2 has an additive effect on cell proliferation (Fig. 5a and S4b) and maintaining PM fluidity (Fig. 5b). These findings confirm that BLTP2 plays a significant role in maintaining PM fluidity in human cells and indicate that some cell types have two independent pathways of regulating PM fluidity, both of which have PE-dependent mechanisms.

**Figure 5.**
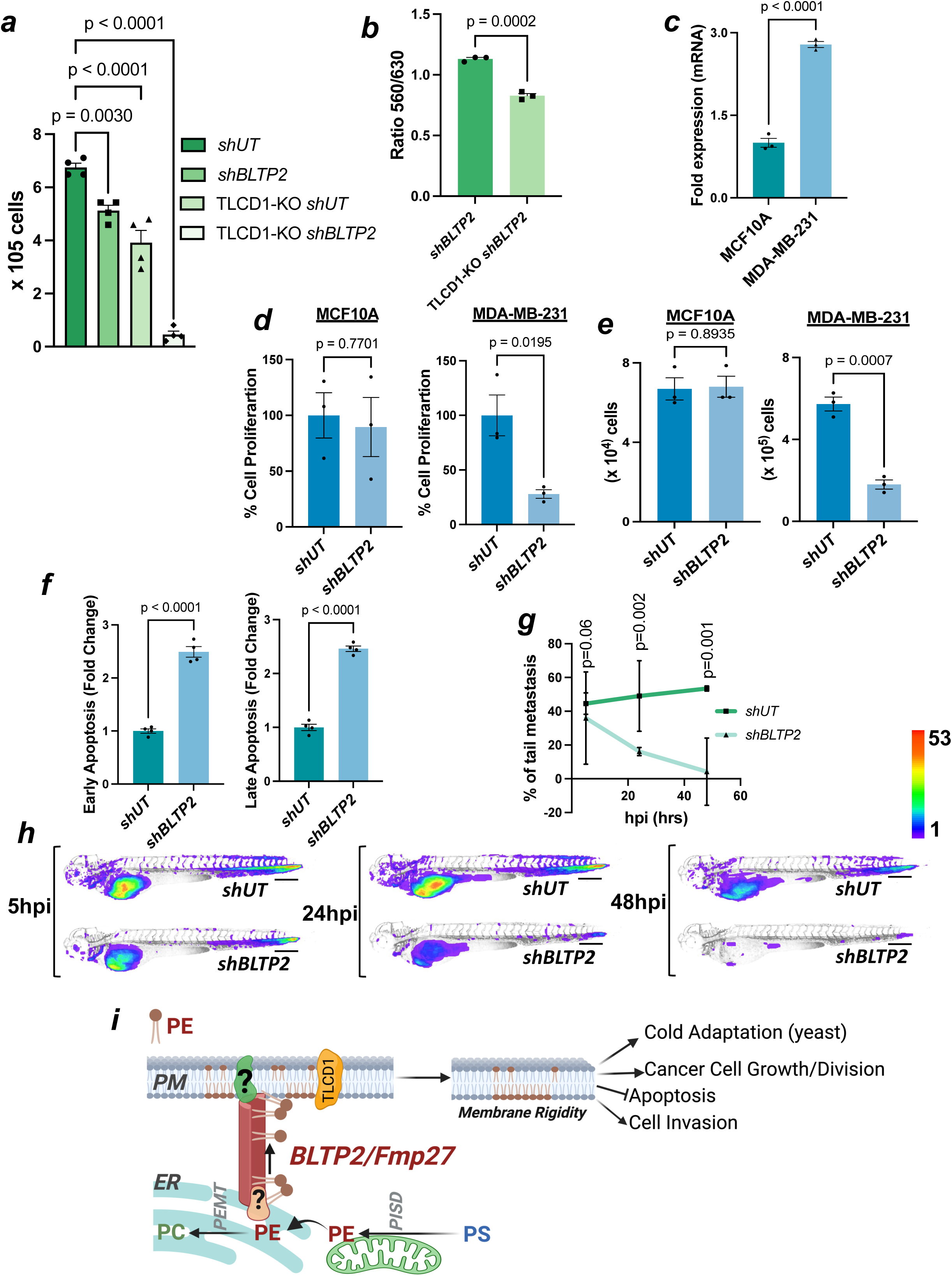
BLTP2 regulates the survival and metastasis of MDA-MB-231 cells and synergizes with TLCD1 to control cell growth and PM fluidity. a. Automated cell counting of HeLa strains grown with DMEM containing complete FBS; mean±SEM (n=4); p-values from one-way ANOVA. b. Ratio of NR12S fluorescence at 560nm and 630nm of HeLa strains grown in DMEM medium with complete FBS and then shifted to DMEM with dFBS for 4 hrs; mean±SEM (n=3); p-values from unpaired t-tests. c. qRT-PCR showing the relative abundance of BLTP2 mRNA in MDA-MB-231 (TNBC) cells compared to MCF10A (non-cancerous) cells; mean±SEM (n=3); p-values calculated from unpaired t-test are indicated. d. e. WST-1 assay (d) and automated cell count assay (e) comparing proliferation of BLTP2-silenced (*shBLTP2*) to control (*shUT*) MCF10A cells (left panels) and MDA-MB-231 cells (right panels); mean±SEM (n=3); p-values from unpaired t-tests. f. Fold difference in the percentage of cells undergoing early (left) and late (right) stages of apoptosis. Apoptotic cells were determined by Annexin-V and propidium iodide dual staining and flow cytometry; mean±SEM (n=4); p-values calculated from unpaired t-tests. g. Percent of xenografted larval zebrafish with tail metastasized MDA-MB-231 expressing *F-tractin* and treated with *shUT* (control) or *shBLTP2* (BLTP2-depleted). Fishes were imaged at 5, 24, and 48 hpi of MDA-MB-231 cells into the yolk. Points indicate mean % of tail metastasized fishes from two independent experiments with shBLTP2 cells (n_1_=74 and n_2_=63 larvae) and shUT cells (n_1_=75 and n_2_=80 larvae); error bars indicate a 95% confidence interval; p-values for untailed t-tests for each time point. h. Quantitative dissemination maps of zebrafish xenografts of MDA-MB-231 expressing *F-tractin* and treated with *shUT* (control) or *shBLTP2* (BLTP2-depleted) using a color-coded scale bar of 1-53 for the number of cells at each location. Scale bars = 500µm. i. Model for the cellular function of BLTP2/Fmp27. The proteins move PE to the PM to regulate PM fluidity and, in humans, work in parallel with TLCD1 PM fluidity-regulating pathway. The scheme is drawn using biorender.com.

To further understand the function of BLTP2 in human cancer cells, we investigated its reported role in the aggressiveness of the triple-negative breast cancer (TNBC) cell line MDA-MB-231^10^. BLTP2 expression is 3-fold higher in these cells than in the non-cancerous breast epithelial cell line MCF10A (Fig. 5c). Acute depletion of BLTP2 (shBLTP2; Fig. S5a) significantly reduced the proliferation rate of MDA-MB-231 cells but not control MCF10A cells (Fig. 5d,e). In addition, the depletion of BLTP2 significantly increased MDA-MB-231 cell apoptosis (Fig. 5f), indicating BLTP2 is a pro-survival protein in TNBC cells. We next tested whether BLTP2 controls the invasiveness of MDA-MB-231 cells in a previously established zebrafish xenograft model^34, 35^. MDA-MB-231 cells stably expressing the actin filament reporter *Ftractin-EGFP* and grown in a medium with FBS were treated with shBLTP2 or shUT and xenografted into larval zebrafish (Fig. S5b). This allowed a quantitative assessment of how these two cell types survive and disseminate *in situ*. Quantifying the survival of micro-metastasized MDA-MB-231 cells in the zebrafish tail, away from the injection site, revealed their survival was significantly reduced after knockdown of BTLP2, particularly at 24 hours and 48 hours post injection (hpi) (Fig. 5g). BTLP2 depletion also significantly affects dissemination; quantitative dissemination maps reveal *shBTLP2* MDA-MB-231 cells have a significantly reduced ability to spread throughout the zebrafish larva (Fig. 5h). Cell migration to the intersegmental vessels or the brain tissue was strongly impaired in BLTP2-depleted MDA-MB-231 cells (Fig. 5h). However, *shBTLP2* cells still spread to the caudal hematopoietic tissue and the gills, perhaps because of passive migration driven by blood flow. Interestingly, migration of BLTP2-depleted MDA-MB-231 cells into upper inter-segmental veins (ISVs) from the caudal hematopoietic tissue is significantly impaired (Fig. S5c). These findings demonstrate that BLTP2-depletion not only hinders TNBC cell proliferation but also significantly impacts its migration in the zebrafish xenografted model. Further evidence was provided by assessing actin ruffling at cell protrusions, a hallmark of cancer cell migration and invasiveness^36^. Imaging the xenografted MDA-MB-231 cells on a high-resolution axially swept light-sheet microscope revealed *shUT*-treated cells formed abundant actin ruffles, but ruffling was reduced in *shBTLP2* cells (Fig. S5d)^37^. We propose that depletion of BLTP2 affects PM fluidity, which alters actin polymerization at the cell surface and reduces migration and invasiveness.

Together, our findings indicate that BLTP2 is part of a conserved pathway that regulates PM fluidity by increasing the amount of PE in the PM. Because the protein is part of an LTP family and localizes to ER-PM contacts, it may directly transport PE from the ER to the PM. What drives the net movement of PE to the PM remains to be determined. However, it should be noted that what determines transport directionality is not known for any member of the Vps13-like family of LTPs. Our finding that BLTP2 and TLCD1 both regulate PM fluidity independently in a PE-dependent manner suggests cells may have more than one mechanism for ensuring there is sufficient PE in the PM. Why BLTP2 is particularly important for the proliferation of several breast cancer cell lines and the aggressiveness of MDA-MB-231 and perhaps other breast cancer cell types is an open question. Our results suggest BLTP2 is pro-tumoral and regulates invasiveness, presumably by controlling the PM fluidity of some cancer cells. These cells may have defects in the endogenous production of EtN, rendering them particularly dependent on exogenous EtN or BLTP2 to maintain PM fluidity. Whether BLTP2 controls tumorigenesis in a PE– and PM fluidity-dependent manner and whether BLTP2 is a targetable protein in aggressive breast cancer treatment will be determined in future studies.

## Acknowledgements

We thank Dr. Grzegorz Piszczek, the director of the biophysics core at NHLBI/NIH. The lipid analysis was conducted on the LIPANG (Lipid analysis in Grenoble) platform hosted by the LPCV, Université Grenoble Alpes, and supported by the Rhône-Alpes Region, the fonds FEDER, and GRAL, financed within the University Grenoble Alpes graduate school (Ecoles Universitaires de Recherche) CBH-EUR-GS (ANR-17-EURE-0003). We thank Dr. Avik Dutta, NICHD/NIH, and Dr. Hijai Regina Shin, UC Berkeley for valuable suggestions on experimental methods. We thank Dr. Andy Golden, Dr. Daniel Masison, Dr. John Hanover, and Dr. Orna Cohen-Fix laboratories for their support. We thank the Animal Resource Center (ARC) and Dr. Gaudenz Danuser’s lab for the zebrafish facility. We thank the members of the Prinz lab for their helpful comments. The research was supported by the NIDDK Intramural Research Funding and by the UT Southwestern Medical Center. The zebrafish research was partly supported by NIH R35GM133522 and U54CA268072 to Dr. Reto Fiolka and computational resources provided by the BioHPC supercomputing facility located in the Lyda Hill Department of Bioinformatics, UT Southwestern Medical Center.

---

**Figure S1.**
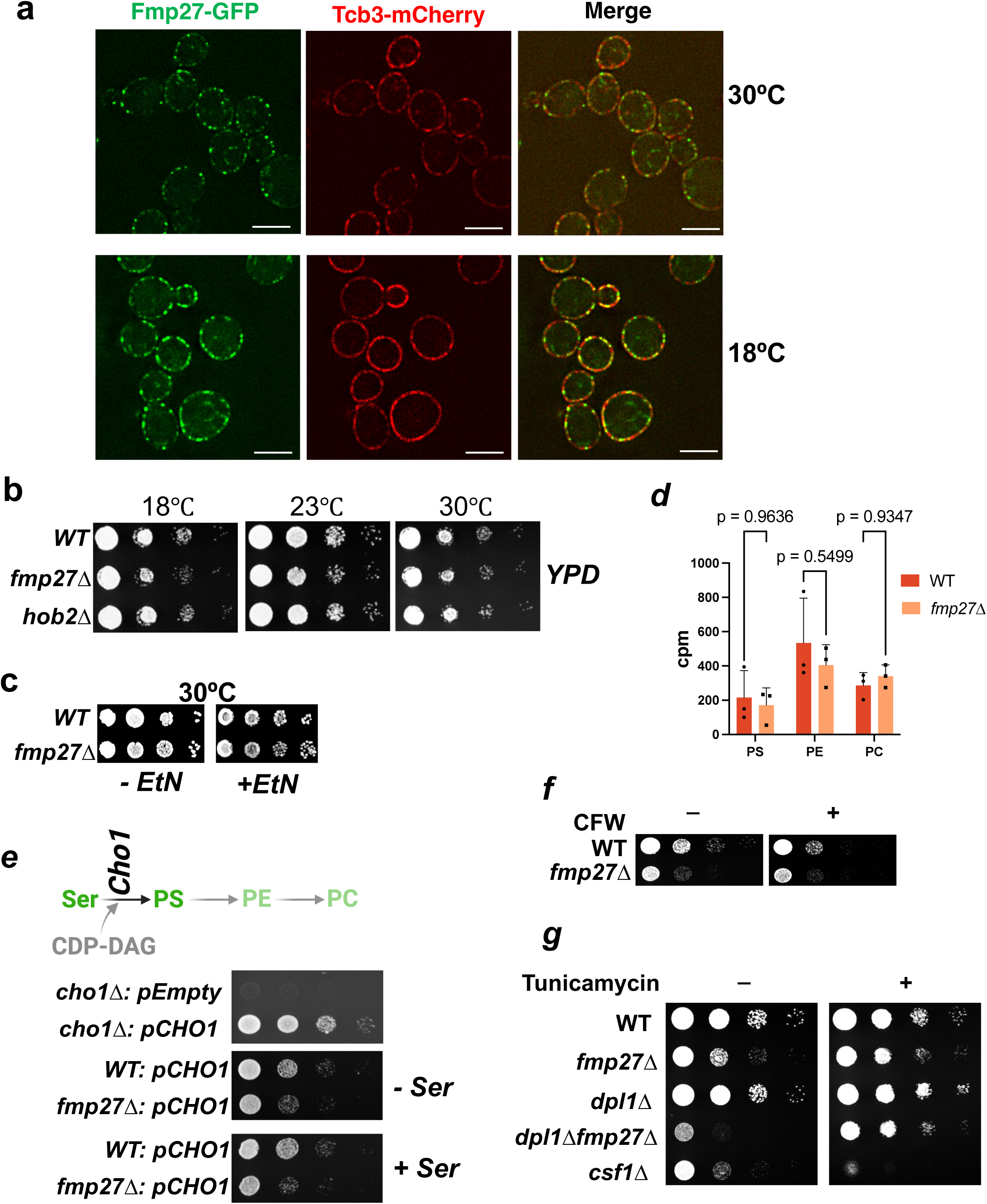
Fmp27 localizes to ER-PM contact sites and cells lacking Fmp27 do not have defects producing PLs by the CDP-DAG pathway. a. Images of yeast cells expressing endogenously tagged Fmp27-GFP and Tcb3-mCherry and grown at 30°C (top) or 18°C (bottom). Scale bars = 5µm. b. Serial dilution of yeast strains spotted on YPD plates and incubated for 3 days (30°C), 4 days (23°C) or 5 days (18°C). c. Serial dilution of yeast strains spotted on SC plates with or without 4mM EtN and incubated at 30°C for 3 days. d. Values used to calculate results shown in Fig. 1j; counts per minute (cpm) of [^3^H] in each PL; mean±SEM (n=3); p-values from unpaired t-tests. e. Scheme showing how PS synthase (Cho1) uses serine (Ser) and CDP-DAG to form PS, which gets converted to PE and then PC (top). The bottom panels show a serial dilution of yeast strains spotted on SC media plates and grown at 18°C for 3 days. The strains contained an empty plasmid (pEmpty) or a plasmid expressing the *CHO1* gene under a GPD promoter (pCHO1). Where indicated, the media was supplemented with 4mM serine. f. Serial dilution of yeast strains spotted on SC medium without (left) or with (right) 10µg/ml calcofluor white (CFW). Plates were grown at 18°C for 3 days. g. Serial dilution of yeast strains spotted on SC medium without (left) or with (right) 0.5µg/ml tunicamycin. Plates were grown at 23°C for 3 days.

**Figure S2.**
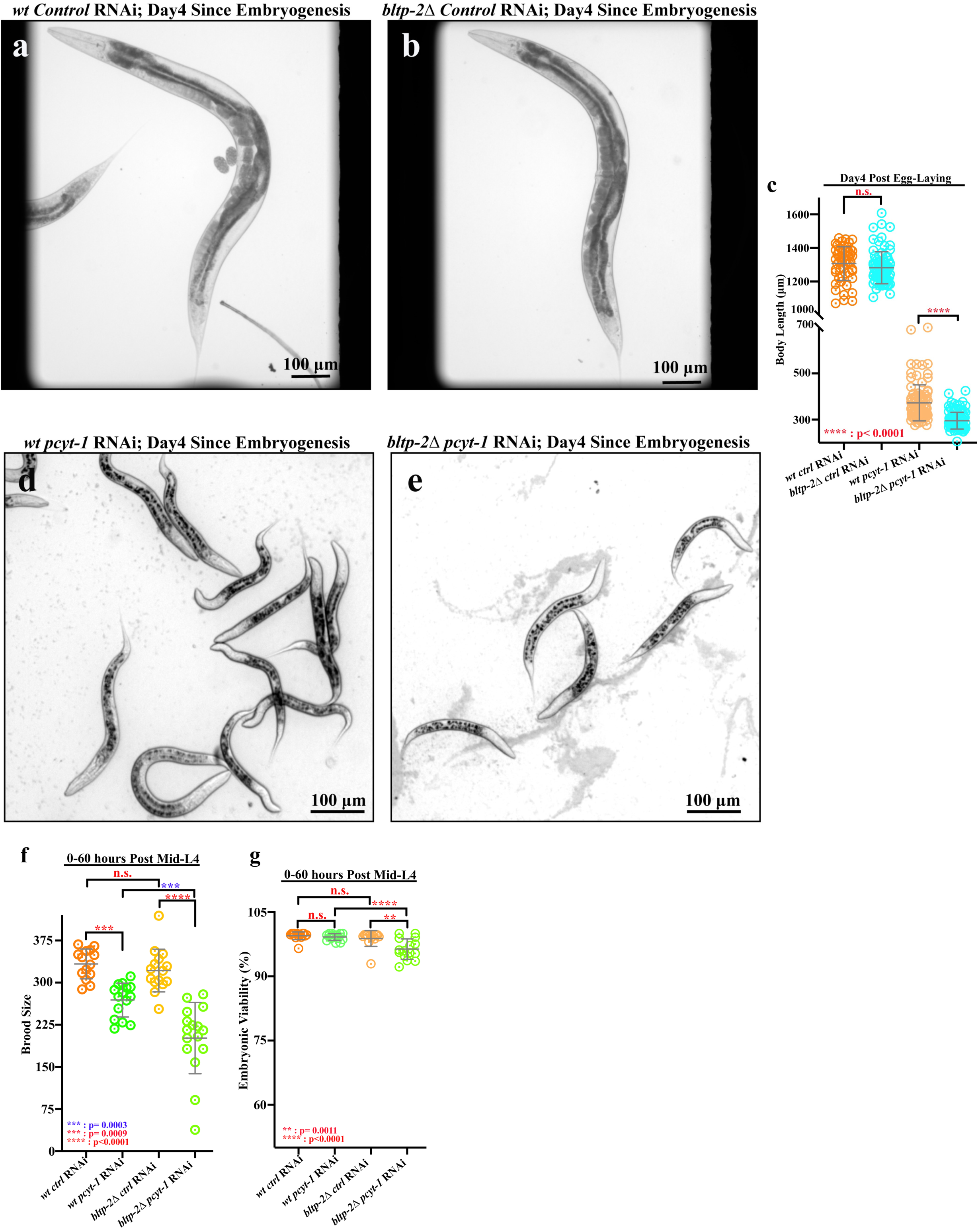
The *C. elegans* gene *bltp-2* coordinates with the Kennedy pathway to maintain growth and physiology. a-b. Representative images of wildtype and *bltp-2Δ* animals treated with control RNAi (a) or *bltp-2* RNAi on day 4 post egg-laying. c. Quantification of the results shown in a. and b; p-values: ****, p<0.0001 (unpaired t-tests). d-e. Depletion of *pcyt-1* by RNAi causes larval arrest in wildtype and *bltp-2Δ* mutant animals. The body length is synergistically smaller in the *bltp-2Δ* mutant compared to the wild-type control. f. A smaller brood size was observed in both wildtype and *bltp-2Δ* when depleted of *pcyt-1* by RNAi. The *pcyt-1* RNAi synergistically reduced the brood size in *bltp-2Δ* compared to wildtype. g. Embryonic viability was reduced in *bltp-2Δ* after depletion of *pcyt-1* by RNAi. P-values: **: p= 0.0011, ***: p=0.0009, ****, p<0.0001 (unpaired t-tests).

**Figure S3.**
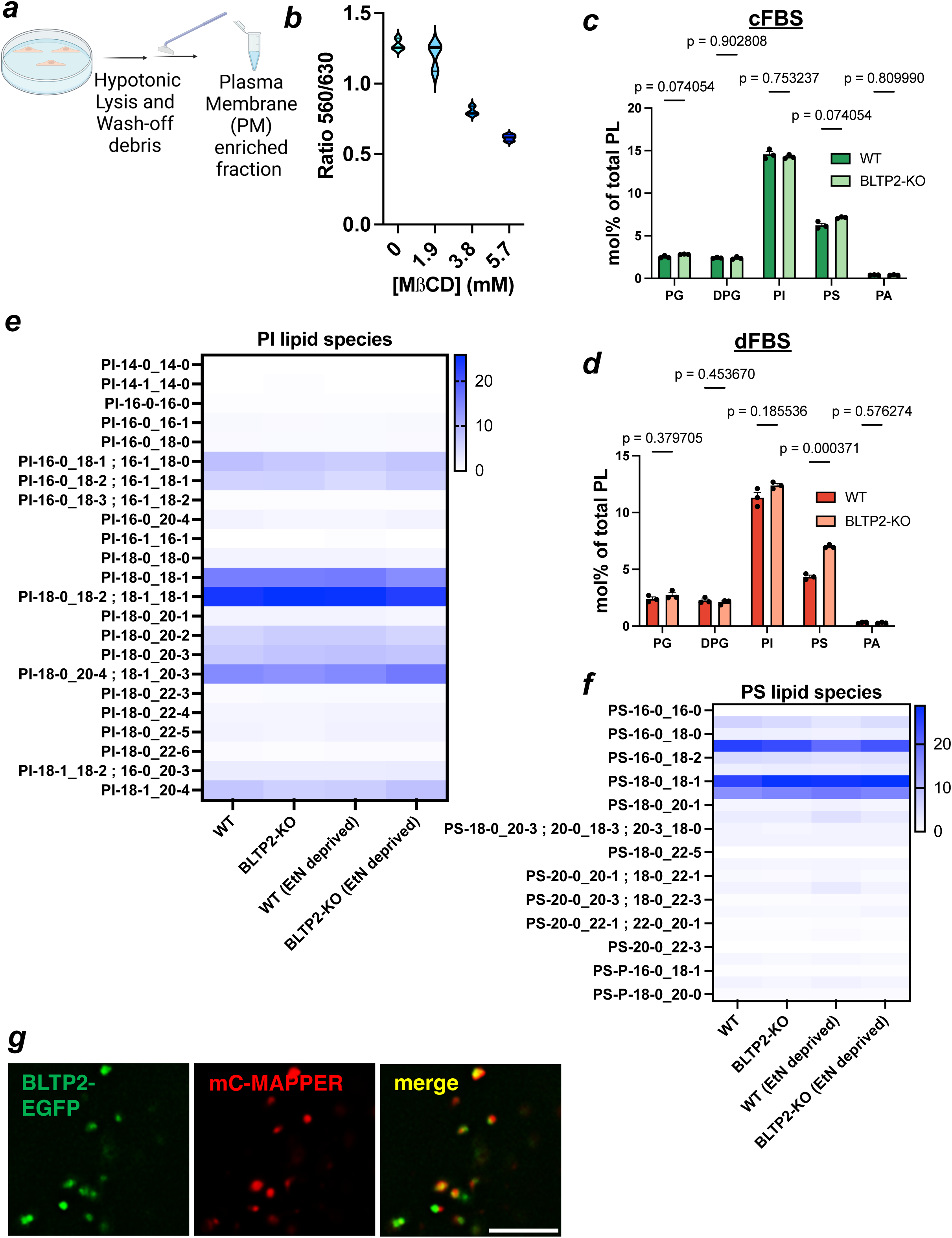
Controls for results shown in Fig. 4. a. Scheme demonstrating the method of isolating a PM-enriched fraction from HeLa cells, drawn using biorender.com. b. Violin plot of the ratiometric fluorescence intensity of NR12S (Y-axis) and concentration of [MßCD] (X-axis). Values from 4 independent experiments are shown. c. Relative abundance of PL species (except PE and PC), determined by LE-MS/MS, from HeLa cells grown in DMEM with complete FBS (cFBS). Values from 3 independent replicates are shown. d. Relative abundance of PL species (except PE and PC), determined by LE-MS/MS, from HeLa cells grown in DMEM with dialyzed FBS (dFBS, which is EtN-deprived)). Values from 3 independent replicates are shown e-f. Results from experiment shown in Fig. 4g for PI (e) and PS (f) species. g. Images of U2OS cells transiently expressing the ER-PM marker mCherry-MAPPER ^38^ and BLTP2-EGFP.

**Figure S4.**
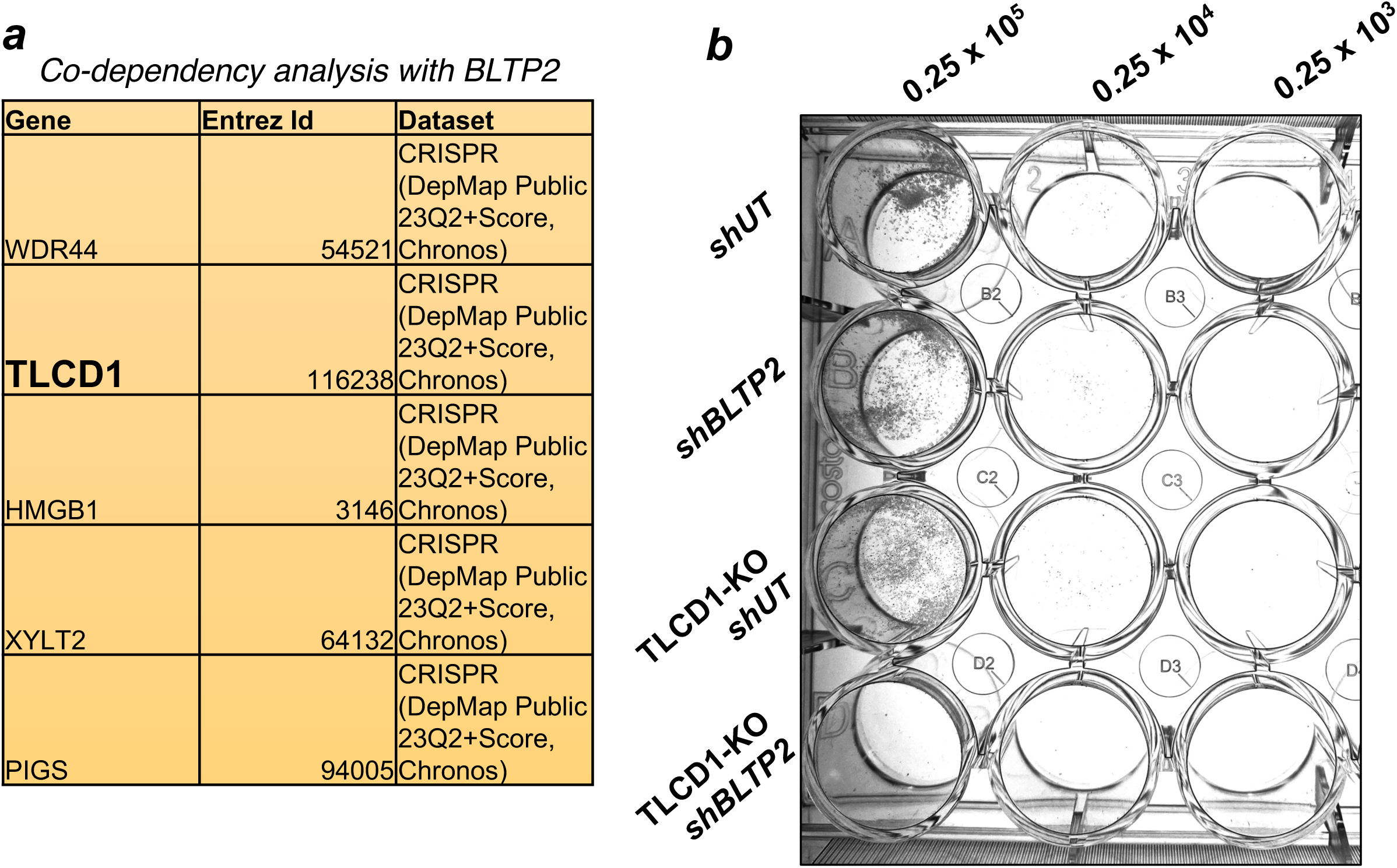
Supplemental finding related to Fig. 5a. a. Table of top 5 co-dependencies of the BLTP2 gene from the CRISPR dataset of DepMap^13, 14^ b. Crystal violet-stained colony forming units of HeLa cells with the indicated genotypes plated at the indicated cell number.

**Figure S5.**
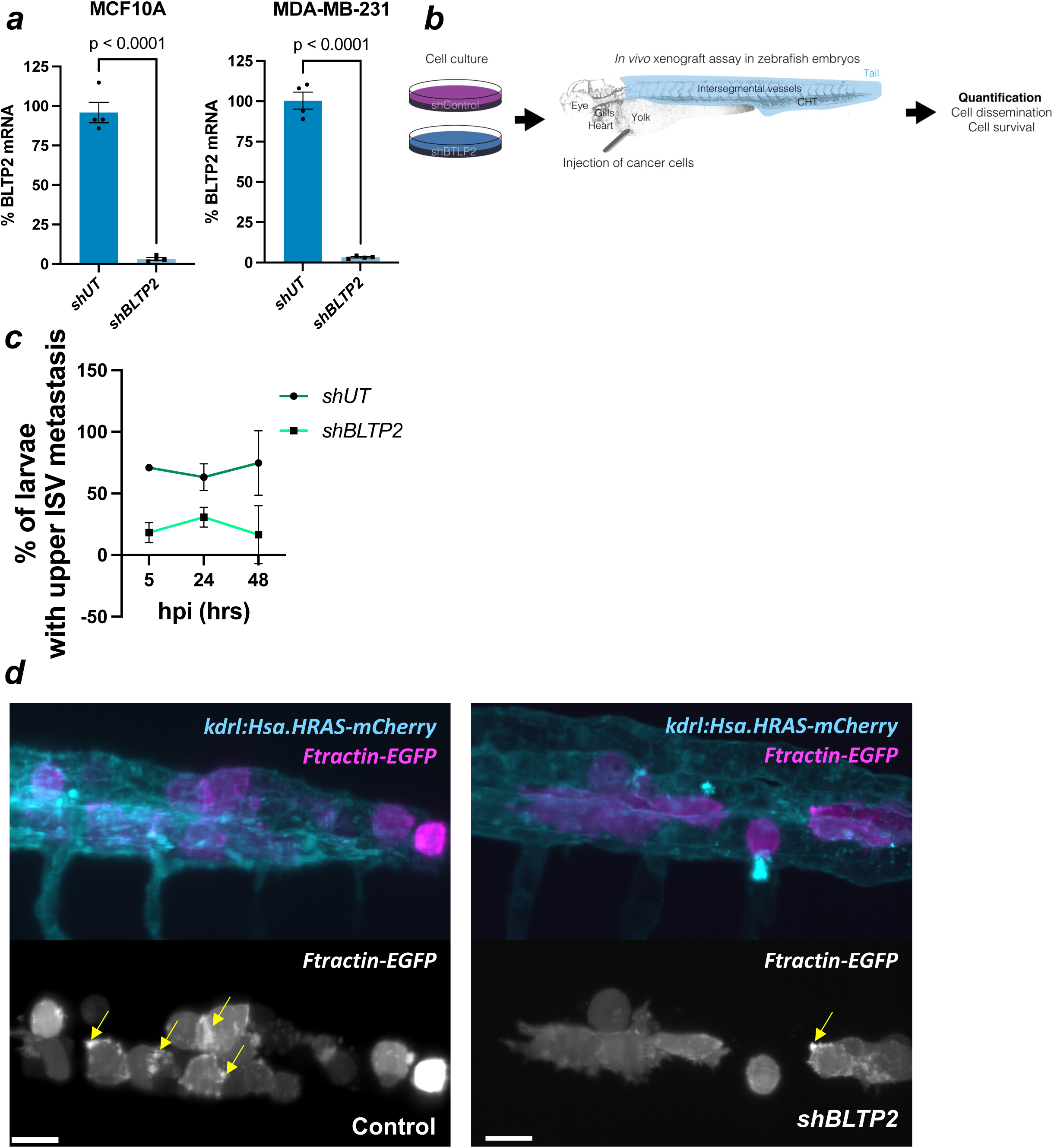
BLTP2 controls actin ruffling in MDA-MB-231 *F-tractin* cells. a. qRT-PCR verifying the knockdown of BLTP2 mRNA when silenced with a shBLTP2 cocktail in MCF10A (left) and MDA-MB-231 (right) cells. b. Schematic of *in vivo* xenograft assay with MDA-MB-231 *F-tractin* cells in zebrafish larvae. c. Top panel shows localization of MDA-MB-231 cells stably expressing the cytoplasmic actin filament reporter *Ftractin-EGFP* (magenta) in zebrafish blood vessels (*Tg(kdrl:Hsa.HRAS-mCherry),* cyan); bottom panel shows actin ruffling (indicated by yellow arrows) at PM protrusion sites in control (left) and shBLTP2 cocktail (right) treated MDA-MB-231 cells. Maximum intensity projections of volumetric data acquired with axially swept light-sheet microscopy are shown. Scale bars = 25µm.

## Materials and Methods

### Yeast strains and culture

Yeast strains used in the study are listed in Table 1. The oligonucleotides used in constructing and verifying yeast strains are mentioned in Table 3. *S. cerevisiae* was grown in either SC medium or YPD medium which was prepared as described in ^4^. When indicated, 4mM Ethanolamine (EtN), Choline (Cho), and Propanolamine (PpN) were added to SC medium. Inositol (Ino) depleted medium was made by adding yeast nitrogen base without inositol (US Biological Life Sciences) and supplemented with 0.1M inositol when indicated. For a serial dilution-based growth assay, yeast cultures were grown to a mid-log phase and spotted in plates containing the indicated medium by a previously published method ^4^.

**Table 1.**
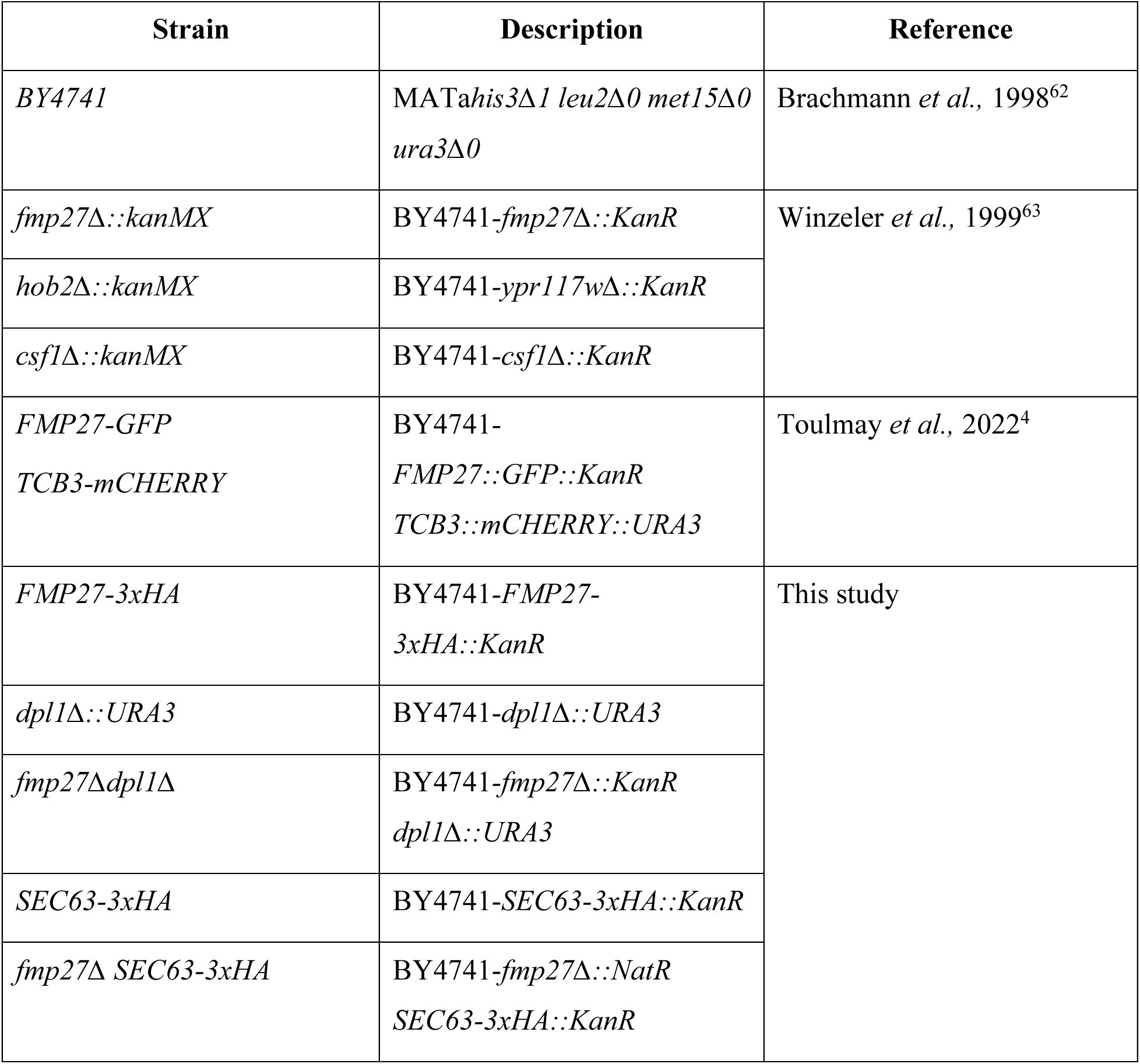
*S. cerevisiae* strains used in this study.

### Mammalian cell culture

HEK293T (Takara Cat. No. 632180), HeLa (ATCC CRM-CCL-2), MDA-MB-231; *F-Tractin-EGFP* cells were grown on DMEM (Gibco Ref. 11965-092) supplemented with 10% fetal bovine serum, penicillin, and streptomycin at 37°C with 5% CO_2_. MDA-MB-231(CRM-HTB-26) cells were grown in Leibovit’s L-15 medium supplemented with 10% fetal bovine serum, penicillin, and streptomycin at 37°C.MCF10A (ATCC CRL-10317) cells were grown in mammary epithelial cell growth medium (Cell Applications Cat. No. 815-500) at 37°C with 5% CO_2_. U2OS cells were grown in McCoy’s 5A Medium (Gibco 16600-082) with 10% fetal bovine serum, penicillin, and streptomycin at 37°C with 5% CO_2_. For experiments without ethanolamine supplementation, DMEM with 25mM HEPES (Gibco Cat. No. 12430-054) supplemented with 10% dialyzed FBS (Gibco A33820-01) was used and 100µM of ethanolamine, choline, and propanolamine was added to the medium.

### Imaging of yeast and mammalian cells

Yeast cells containing genomically tagged *FMP27-GFP* and *TCB3-mCHERRY* were grown to log phase at 30°C. Then subcultures at or below 0.1 OD_600nm_ units were started at indicated temperatures and grown up to an OD_600nm_ = 1. Micrographs of live cells were taken and images were processed by previously used methods^4^.

U2OS cells were first knocked down for endogenous BLTP2 using shRNA (Sigma TRCN0000128410) that targets only the 3’UTR of the BLTP2 mRNA. These cells were then transfected with pcDNA3.1 (+)BLTP2-EGFP. They were also co-transfected with a plasmid containing mCherry-MAPPER, which was kindly gifted by Dr. Jen Liou at UT Southwestern Medical Center ^38^. The cells were imaged 48 hrs after transfection in 4-well Labtech Chambers. Images were processed and merged on FiJi and cropped using Adobe Photoshop 2024.

### Structure prediction and alignment

We first used Deepmind Alphafold to predict the structure of human BLTP2 (Protein KIAA0100; Uniprot Q14667) and yeast Fmp27 (Protein FMP27; Uniprot Q06179). Then, we used UCSF Chimera to perform a structural alignment of HsBLTP2 and ScFmp27 using the Matchmaker plugin.

### BLTP2 score analysis

Publicly available data from DepMap portal of the Broad Institute was used. We plotted the effect of BLTP2-CRISPR deletion [KIAA0100 Gene Effect (Chronos) CRISPR (DepMap Public 23Q2+Score, Chronos)] against the expression level of BLTP2 [KIAA0100 log2(TPM+1) Expression Public 23Q2] available from 1020 cancer cell lines. We highlighted three metastatic breast cancer cell lines as red dots.

### PL measurement by High Performance Liquid Chromatography (HPLC)-Evaporative Light Scattering Detector (ELSD)

We measured PE and PC with an Agilent 1260 Infinity II HPLC and Agilent ELSD. Lipids were isolated from yeast or mammalian cells of indicated genotypes by using a methanol-chloroform extraction as previously mentioned ^4^. Lipids were resuspended in 50µl chloroform and injected with 10µl volume into a Zorbax CN 4.6×150mm column set at 50°C. The samples were run for 15 mins each at a column pressure of 50psi. The mobile phase is composed of Solvent A and B at a 7:3 ratio. Solvent A is Hexane and Solvent B is a w/w mixture of 60% Toluene, 40% Methanol, 0.2% Acetic Acid and 0.1% Triethylamine. A continuous nitrogen gas flow r the ELSD detector was used at a rate of 1.6ml/min. The evaporator and nebulizer temperatures for the ELSD were set at 80°C and 50°C respectively. Integrated peak values for PE and PC were obtained from Agilent OpenLAB and a ratio of PE:PC level was conducted manually.

### Radioactive labeling of PL and quantitative measurements

To label PL to a steady state, 50 ml log phase yeast cultures of the indicated genotypes were gown for up to 5 doublings in SC medium at 18°C with 250µCi of [^3^H]-Palmitic acid (American Radiolabeled Chemicals, ART0129A). For pulse labeling with [^3^H]-Serine, 50 ml of yeast cultures of indicated genotypes were grown in SC medium and were labeled with 100µCi of [^3^H]-Serine (American Radiolabeled Chemicals, ART0246) at 18°C for 1hr. Lipids were isolated, separated by thin layer chromatography, and quantified based on previously established methods ^39^ ^40^ ^4^

### Rapid Immunopurification of Endoplasmic Reticulum (ER) membranes

Yeast strains containing 3xHA tag at the genomic locus of *SEC63* were grown to log phase for seeding cultures. Cultures were seeded overnight and grown up to OD_600nm_ = 1. 25 OD_600nm_ units of yeast cells were mixed with 10mM sodium fluoride and 10mM sodium azide. Cells were pelleted by spinning in a swinging bucket rotor for 1900*g* for 5 mins at 4°C. Cells were washed in 0.5 ml KPBS buffer (136mM KCl, 10mM KH2PO4, pH 7.25). Cells were pelleted in a tabletop centrifuge by spinning them at 3381*g* for 3 mins at 4°C. Cells were lysed using 300µl glass beads in 300µl lysis buffer (KPBS containing protease inhibitor cocktail from Roche Cat. No. 11836170001 and 1mM PMSF). Cell lysis was conducted in a Precellys 24 Homogenizer (Bertin Technologies) with 30 sec homogenization cycles for 3 mins and 1 min gaps to prevent overheating. Cell debris was spun down at 400*g* for 5 mins at 4°C. The supernatant was separated and kept in a fresh tube. 600µl of lysis buffer was added to the glass beads, vortexed, spun down and 550µl supernatant was obtained and mixed with the previous supernatant. 100µl of this supernatant was kept for a cellular lipid measurement. The remaining supernatant was added to 100µl of anti-HA antibody magnetic beads (Pierce Cat. No. 88837) that were prewashed in lysis buffer. After rocking at 4°C for 15 mins the beads were washed 3 times in 1ml cold lysis buffer for 5 mins each at 4°C. For Western blotting, the beads were resuspended in 4X LDS sample buffer (Invitrogen Cat. No. NP0007) containing 2% of 2-mercaptoethanol (Thermo scientific Cat. No: 125470100) and heated at 70°C for 10 mins before freezing them at – 80°C. For lipid extraction, beads were resuspended in 1.6ml de-ionized water, 4ml methanol, and 2ml CHCl_3_. Lipid extraction and radiolabeled lipid measurements were conducted using a previously established method^39^

### Isolation of plasma membranes (PM) and lipid measurement

For the isolation of yeast PM, we used a previously established method^24^. We started with mid-log phase yeast cultures grown at the indicated temperatures. 25 O.D. units of yeast cells were mixed with 10mM sodium fluoride and 10mM sodium azide. Cells were pelleted by spinning in a swinging bucket rotor for 1900*g* for 5 mins at 4°C. Cells were washed in 0.5 ml breaking buffer (50mM Tris pH7.5, 1mM EDTA, 1mMPMSF, and protease inhibitor cocktail). Cells were lysed using 300µl glass beads in 300µl breaking buffer. Cell lysis was conducted by hand vortexing at full speed using a Daiger Vortex Genie2 (Cat. No. 3030A) for 7 times and 45 sec each. Cell debris was spun down at 400*g* for 5 mins at 4°C. The supernatant was collected. 300µl breaking buffer was added, vortexed shortly, spun down and the supernatant was collected and pooled. 500µl of supernatant was mixed well with 500µl 76% Renografin. 800µl of this mix was overlayed with 34%, 30%, 26%, and 22% of Renografin and centrifuged at 193911.2*g* using a Beckman SW55Ti swinging bucket rotor in a Beckman ultracentrifuge. After centrifugation 14 fractions were collected from the top (Fig. 2b). For Western blot to examine organelle membrane fractions, samples were prepared in 4X LDS sample buffer containing 2% 2-mercaptoethanol, heated at 70°C for 10 mins. PM from HeLa cells were isolated following a ‘rip off’ method established by a previously published work^32^ (Fig. S5.a)

For lipid extraction, membrane fractions were resuspended in 1.6ml de-ionized water, 4ml methanol, and 2ml CHCl_3_. Lipid extractions were conducted as mentioned previously. PE and PC lipids were measured by using an HPLC-ELSD detection method as previously described in this section.

### Western blotting

Protein samples were loaded on NuPAGE pre-cast polyacrylamide gels (Thermofisher Scientific) for SDS-PAGE electrophoresis. Pageruler (and Pageruler plus) prestained protein ladders (Thermofisher scientific Cat. No. 26616 and 26619) were used as protein molecular weight standards. Separated proteins were transferred to 0.2µm and 0.45µm nitrocellulose membranes (Biorad Cat. no. 1620115 and 1620112) for transferring proteins below and above 180kDa, respectively. The membranes containing the transferred proteins were blocked in a blocking buffer (5% non-fat milk in 0.1%TBST). Membranes were probed with the appropriate primary antibodies following the dilutions that are instructed in the manufacturer’s protocol (refer to Table 2) in blocking buffer containing either 5% non-fat milk or 5% bovine serum albumin in 0.1%TBST overnight at 4°C. On the following day, the membranes were washed in 0.1%TBST, 3 times for 5 mins each. Membranes were incubated with 1:8000 dilutions of LI-COR secondary antibodies (refer to Table 2) in a blocking buffer for 1 hour at room temperature. Membranes were again washed in 0.1%TBST, 3 times for 5 mins each. The membranes were imaged directly in an Odyssey LICOR imager.

**Table 2.** Antibodies used in this study.

| Antibodies | Source |
| --- | --- |
| anti-HA | Cell signaling technology Cat. no. C29F4 |
| anti-Por1 | Thermofisher scientific Cat. No. 459500 |
| anti-Pma1 | Gift from the Lab of Dr. Amy Chang,<br>University of Michigan, Ann Arbor, USA |
| anti-Dpm1 | Thermofisher scientific Cat. No. A6429 |
| anti-Vph1 | Abcam 10D7A7B2 Cat. No. ab113683 |
| anti-Caveolin-1 | Cell signaling technology Cat. no. D46G3 |
| anti-GAPDH | Cell signaling technology Cat. no. 14C10 |
| anti-VDAC | Cell signaling technology Cat. no. D73D12 |
| anti-PDI | Cell signaling technology Cat. no. C81H6 |
| IRDye 800CW anti-Rabbit IgG | LI-COR 926-32211 |
| IRDye 680RD anti-mouse IgG | LI-COR 926-68070 |

**Table 3.**
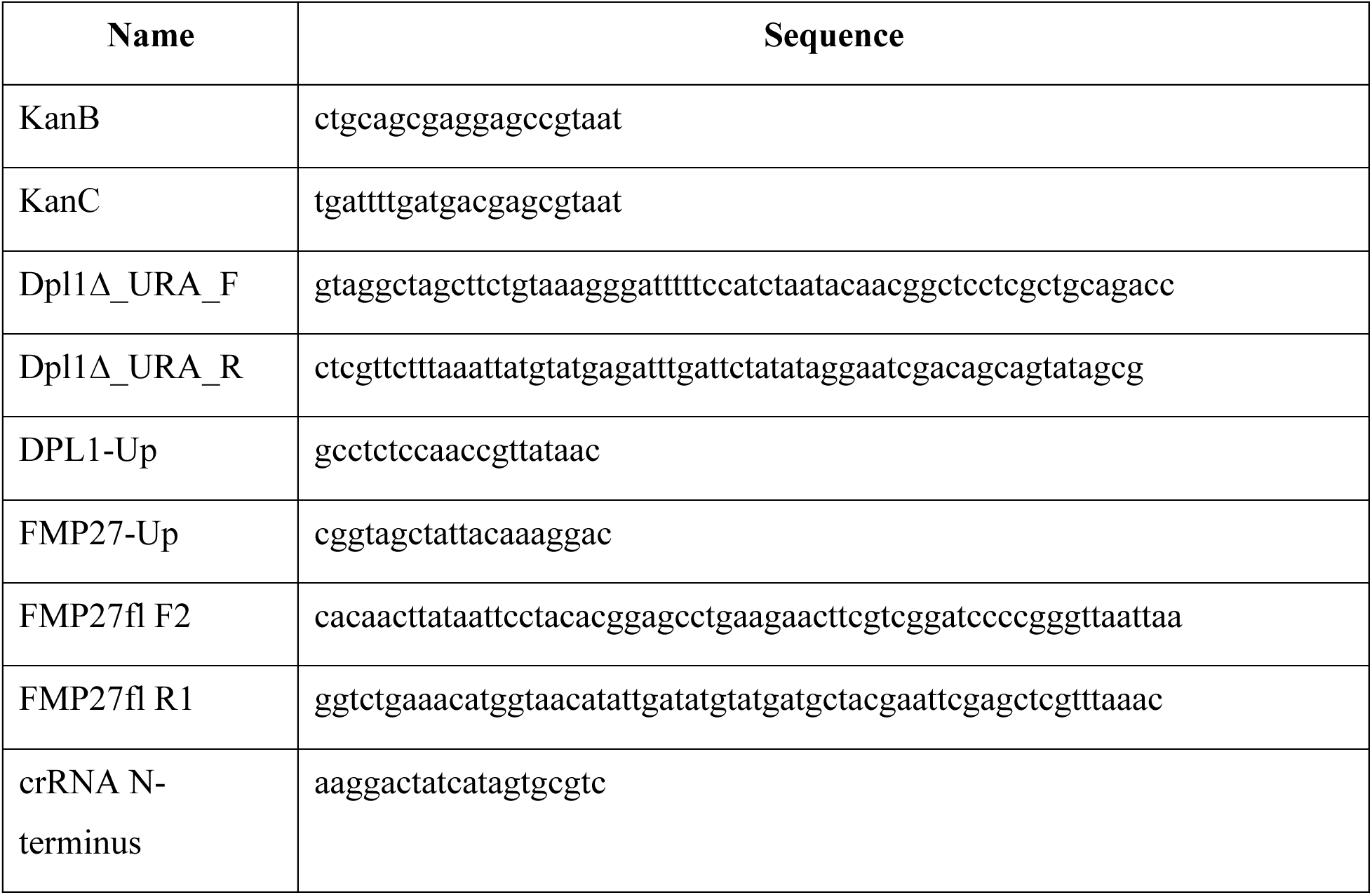
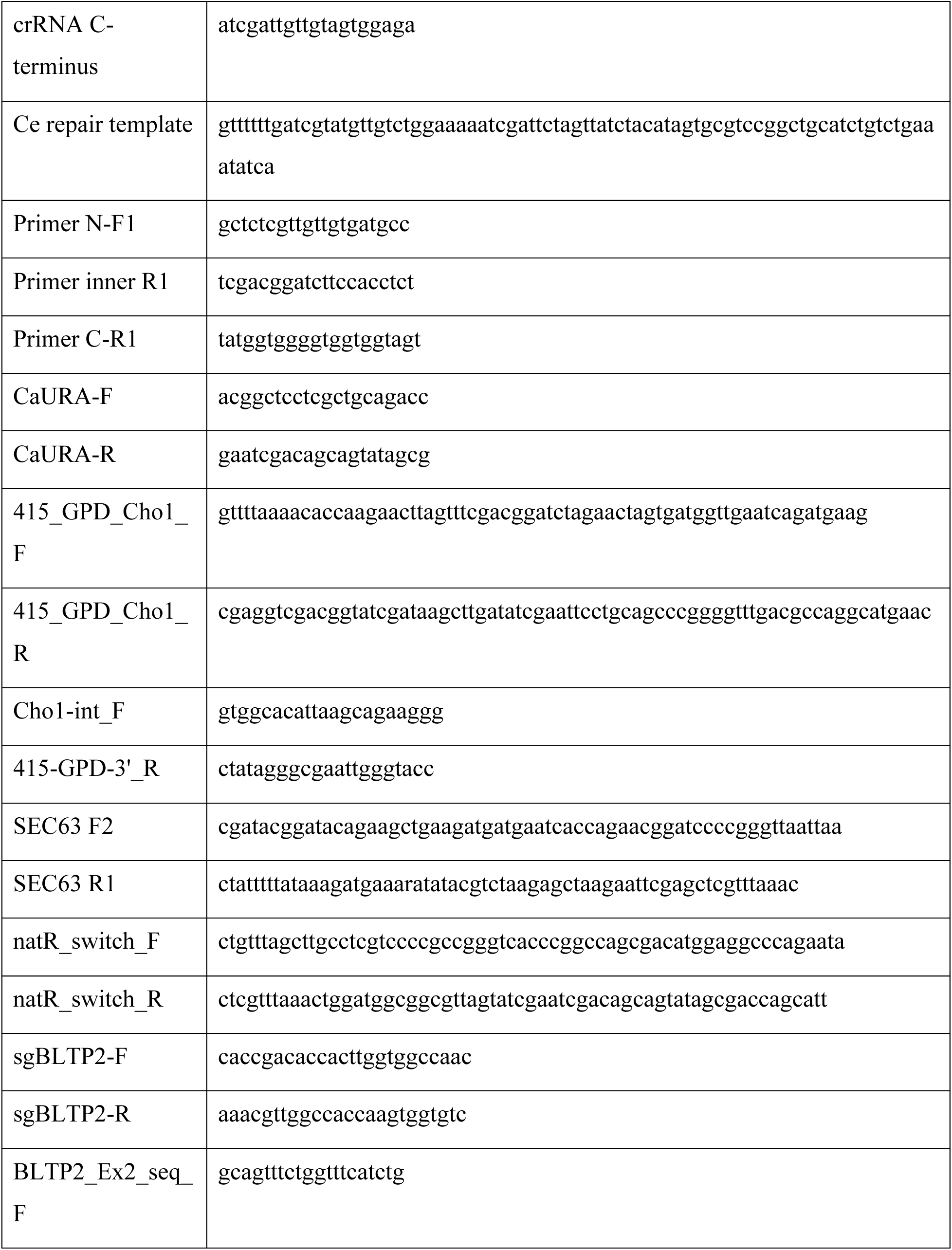
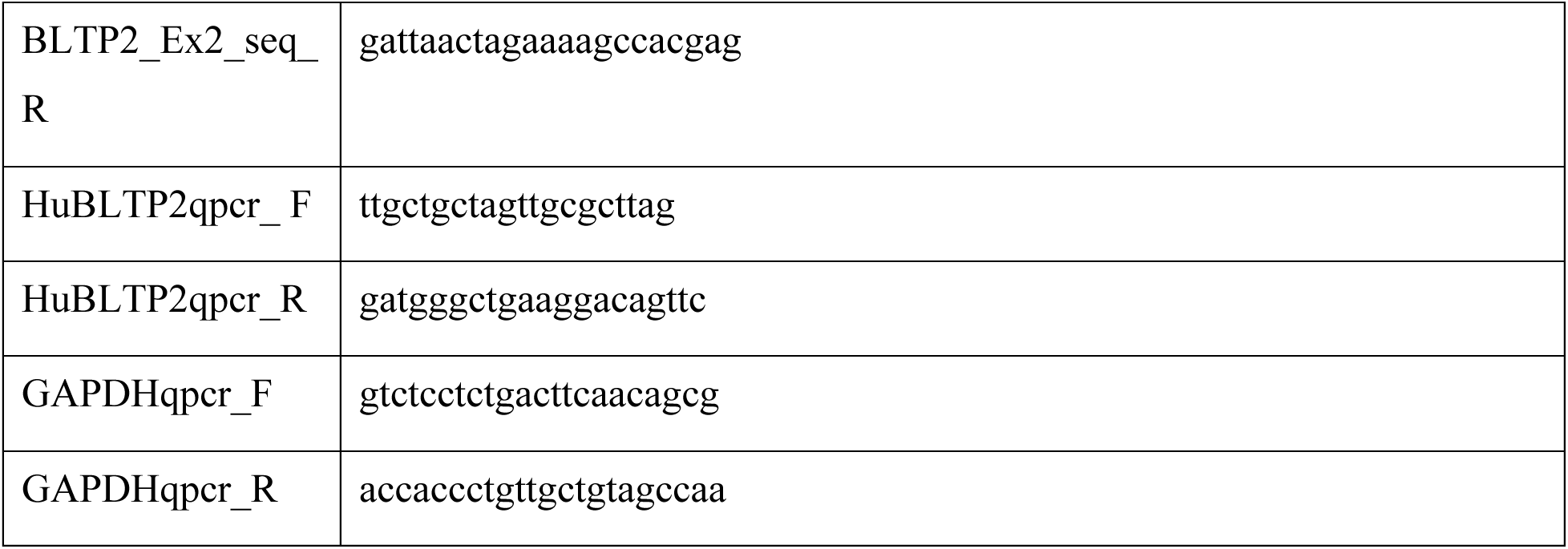
Oligonucleotides used in this study.

### Measurement of plasma membrane fluidity

Yeast cells of the indicated genotype were grown to mid-log phase at the indicated temperatures. To measure the fluidity of yeast PM, we used 1-(4-Trimethylammoniumphenyl)-6-Phenyl-1,3,5-Hexatriene p-Toluenesulfonate (TMA-DPH; Thermofisher Scientific Cat. No. T204). Yeast cells were stained with 0.5µM TMA-DPH, as previously conducted in^25^. The steady-state fluorescence anisotropy of TMA-DPH at the indicated growth temperatures was measured using a Horiba spectrofluorometer at the Biophysics core of the National Heart, Lung, and Blood Institute (NHLBI/NIH). The steady-state anisotropy (*r*_s_) was calculated using a previously established method^25^.

The PM fluidity of mammalian cells was measured using 3-((3-((9-(Diethylamino)-5-oxo-5H-benzo[a]phenoxazin-2-yl)oxy)propyl)(dodecyl)(methyl)ammonium)propane-1-sulfonate (NR12S; Tocris Bioscience Cat. No. 7509) using a method as exactly mentioned in Wendy S. Smith, Sopsamorn U. Flavell, David J. Flavell, The Simon Flavell Leukaemia Research Laboratory, Southampton General Hospital, Southampton, 02/2017 (BMG LABTECH AN302). The NR12S-stained cells were excited at 520nm and fluorescence emissions were measured at 560nm and 630nm using a BMG Omega plate reader. A ratio of fluorescence intensity at 560nm to the intensity at 630nm was manually calculated.

### Production of lentivirus and infection

Lentiviruses were produced and transduced using a previously established method^41^. Precisely, lentiviral plasmid (pLKO.1) containing shRNA expressing sequences were co-transfected with packaging vectors psPAX2 and pMD2G into HEK293T cells. Media containing the virus was collected after 48hrs filtered through a 0.45µm PES low binding filter. This was then concentrated using Lenti-X concentrator (Clontech Cat. No. 631231). 5 shBLTP2 viral pellets were mixed into an shRNA cocktail in 1ml complete DMEM medium. Control shUT viral pellet were reconstituted in 0.2ml ml complete DMEM for an equal dosage of control shUT virus in our experiments. Finally, the reconstituted lentiviruses were stored at –80°C for future use. shRNAs depleting BLTP2 (TRCN0000128930, TRCN0000129099, TRCN0000130217, TRCN0000128410, TRCN0000129809) was purchased from Millipore-Sigma as bacterial glycerol stocks. shUT containing an untargeted shRNA was used as previously published in our work^4^. Cells were infected with lentiviruses and 8ug/ml of Polybrene (Millipore-Sigma TR-1003-G). Growth medium was changed to fresh medium containing puromycin for selection. Experiments were conducted with puromycin-selected cells within 5-7 days post-infection.

### qRT-PCR

We used qRT-PCR to verify the knockdown of BLTP2 using a previously established method^4^. For reverse transcription PCR, a qScript cDNA supermix (Quantabio Cat. No. 95048-025) was used and the manufacturer-recommended protocol was used. For qPCR, PowerTrack SYBR green master mix (ThermoFisher Scientific Cat. No. A46012) and a manufacturer-recommended protocol for a 20µl reaction were used. qPCR was conducted using a Biorad C1000 Touch Thermal Cycler (CFX96 Real-Time system). The oligonucleotides used for qRT-PCR of BLTP2 mRNA are HuBLTP2qpcr_ F and HuBLTP2qpcr_R. GAPDH mRNA was used as internal control and qPCR was conducted using the oligonucleotides GAPDHqpcr_F and GAPDHqpcr_R.

### Generation of CRISPR KO cell lines

BLTP2 was knocked out in HeLa cells using the pSpCas9(BB)-2A-Puro plasmid^42^ and sgRNA targeting Exon2. sgRNA was ordered as oligonucleotides (sgBLTP2-F and sgBLTP2-R) and cloned using a BbsI restriction enzyme digested vector according to the method mentioned by the Feng Zhang lab at the MIT in Addgene (Addgene #62988)^42^. We followed a previously established method ^43^ to obtain a single clone of BLTP2 knock-out cell line (BLTP2-KO) and verify BLTP2-KO clones by PCR using the oligonucleotides BLTP2_Ex2_seq_F and BLTP2_Ex2_seq_R followed by sanger sequencing of the purified PCR product. Precisely, cells were transfected with pSpCas9(BB)-2A-Puro plasmid containing the sgRNA with Fugene HD transfection reagent (Promega Cat. No. E2311) following the manufacturer’s protocol. Cells were selected with Puromycin and plated in a 96-well plate to obtain single clones. gDNA from clones was extracted using a gDNa extraction kit (Zymo Research Cat. No. D3024) and PCR was conducted using KAPA HiFi DNA Polymerase (Roche Cat. No. KK2601). The verified BLTP2-KO clone was grown in 10cm tissue culture plates and stored as liquid nitrogen stocks for subsequent experiments. The TLCD1-KO and isogenic control HeLa cell lines were kindly gifted by Dr. Kasparas Petkevicius, MRC Mitochondrial Biology Unit at Cambridge University, UK^7^.

### Cell proliferation assays

Human-derived cell lines of indicated genotype were plated at equal density for a particular experiment. For an automated cell count assay^44^, experiments were conducted in a 6-well plate or a 24-well plate. 10^5^ cells in a 6-well plate were plated for each genotype and grown in the indicated medium. For experiments conducted in 24-well plate, 0.25×10^5^ cells were plated for each genotype and condition. On days 4-5, cells were trypsinized with 0.5-1 ml 0.25% trypsin/EDTA and resuspended in 1-3ml of growth medium. Adding 10µl trypan blue to 30µl of cell suspension, 10µl of the mix was injected in a countess chamber slide (Invitrogen), and cell number/ml was counted in Countess 3 (Invitrogen by Thermofisher Scientific). Cell numbers were corrected for trypan blue addition and volume of resuspension and presented as histograms.

For colony formation assay crystal violet staining was used. A serial 10-fold dilution of cells of the indicated genotypes was plated. After days 4-5 plates were taken out on ice, and washed 2-times with ice-cold 1X DPBS (Gibco Cat. No. 14190-144). Cells were fixed with ice-cold methanol for 10 minutes. After aspirating methanol from plates, plates were removed from ice and enough 0.5% crystal violet solution was added to cover the bottom of the plate. After incubating at room temperature for 10 minutes crystal violet solution was discarded. The wells were carefully rinsed in deionized water until the blue color was no longer coming off in the rinse. Plates were dried at room temperature overnight. Images were taken either in UVP GelSolo Imager by Analytikjena or using a transilluminator and iPhone 11 camera. WST-1 assays to measure cell proliferation rate (Roche Cat. No. 5015944001) were conducted using the manufacturer’s protocol.

### Annexin-V apoptosis assay

Flow cytometry-based detection of early and late apoptosis of cancer cells was conducted using the Annexin-V apoptosis assay kit (Thermofisher Scientific Cat. No. V13241). Staining of cells with Annexin-V and propidium iodide was conducted according to the manufacturer’s protocol. The stained cells were analyzed by Flow Cytometry using the UT Southwestern flow cytometry core and plotted as histograms using GraphPad Prism.

### LE-MS/MS based lipidomics

Total fatty-acid quantification was performed using gas-chromatography with Flame Ionization Detector (GC-FID) as described in^45^. For LC-MS/MS analyses, the lipid extracts corresponding to 25 nmole of total fatty acids were dissolved in 100 µL of chloroform/methanol [2:1, (v/v)] containing 125 pmole of each internal standard. Internal standards used were PE 18:0-18:0 and DAG 18:0-22:6 from Avanti Polar Lipid and SQDG 16:0-18:0 extracted from spinach thylakoid^46^ and hydrogenated as described in^47^. Lipids were then separated by HPLC and quantified by MS/MS.

The HPLC separation method was adapted from a previously published method^25^. Lipid classes were separated using an Agilent 1200 HPLC system using a 150 mm×3 mm (length × internal diameter) 5 µm diol column (Macherey-Nagel), at 40°C. The mobile phases consisted of hexane/isopropanol/water/ammonium acetate 1M, pH5.3 [625/350/24/1, (v/v/v/v)] (A) and isopropanol/water/ammonium acetate 1M, pH5.3 [850/149/1, (v/v/v)] (B). The injection volume was 20 µL. After 5 min, the percentage of B was increased linearly from 0% to 100% in 30 min and stayed at 100% for 15 min. This elution sequence was followed by a return to 100% A in 5 min and an equilibration step for 20 min with 100% A before the next injection, leading to a total runtime of 70 min. The flow rate of the mobile phase was 200 µL/min. The distinct glycerophospholipid classes were eluted successively as a function of the polar head group.

Mass spectrometric analysis was done on a 6470 triple quadrupole mass spectrometer (Agilent) equipped with a Jet stream electrospray ion source under following settings: Drying gas heater: 260°C, Drying gas flow 13 L/min, Sheath gas heater: 300°C, Sheath gas flow: 11L/min, Nebulizer pressure: 25 psi, Capillary voltage: ± 5000 V, Nozzle voltage ± 1000. Nitrogen was used as collision gas. The quadrupoles Q1 and Q3 were operated at widest and unit resolution respectively. PC and Lyso-PC analysis was carried out in positive ion mode by scanning for precursors of m/z 184 at a collision energy (CE) of 34 eV. SQDG analysis was carried out in negative ion mode by scanning for precursors of m/z –225 at a CE of –56eV. PE, PI, PS, PG, PA, and Lyso-PE measurements were performed in positive ion mode by scanning for neutral losses of 141 Da, 277 Da, 185 Da, 189 Da, 115 Da, and 141 Da at CEs of 20 eV, 12 eV, 20 eV, 16 eV, 16 eV, and 20 eV, respectively. Plasmanyl-ethanolamine (PE-A) and plasmanelyl-ethanolamine (PE-P) measurements were performed in positive ion mode by scanning for *sn-1* ether + C_2_H_8_NO_3_P product at CE of 20 eV ^48^. Quantification was done by multiple reaction monitoring (MRM) with 30 ms dwell time. DAG and TAG species were identified and quantified by MRM as singly charged ions [M+NH_4_]^+^ at a CE of 16 and 22 eV respectively with 30 ms dwell time. CL species were quantified by MRM as singly charged ions [M-H]-at a CE of –45 eV with 50 ms dwell time. Mass spectra were processed by MassHunter Workstation software (Agilent) for identification and quantification of lipids. Lipid amounts (pmole) were corrected for response differences between internal standards and endogenous lipids and by comparison with quality control (QC). QC extract corresponds to a known lipid extract from yeast qualified and quantified by TLC and GC-FID as described by^49^. Analyzed transitions are based on a previously published method ^50^ for yeast and on published methods ^48^ ^51^ ^52^ for mammalian samples.

### Data presentation of lipid species

Mole percentage of measured glycerophospholipid species were measured from triplicate cultures of indicated yeast genotype. The logarithmic (log to the base2) fold change of lipid species from yeast grown at the indicated temperature with or without EtN supplementation was measured using MS Excel. An unpaired t-test was conducted to obtain p-values between conditions and converted to a log value using MS Excel. Volcano plots were plotted using GraphPad Prism 10.

An Unsaturation Index to measure the number of double bonds per glycerophospholipid was used to calculate changes in lipid unsaturation for yeast of indicated genotype, grown at the indicated temperature, with or without EtN supplementation in the medium. This was calculated following a previously described method^53^.

For mammalian glycerophospholipid species analysis, the mole percentage of indicated glycerophospholipid species were measured from triplicate cultures of cells and analyzed using the heat map feature of GraphPad Prism 10.

### Zebrafish husbandry and xenograft sample preparation

Zebrafish husbandry and experiments described here have been approved and conducted under the oversight of UT Southwestern’s Institutional Animal Care and Use Committee (IACUC) under protocol number 101805 to Gaudenz Danuser. Zebrafish adults and embryos were kept at 28.5°C and were handled according to established protocols^34^. To visualize cancer cells in a near-physiological environment *in situ*, a zebrafish (*Danio rerio*) line expressing the vascular marker *Tg(kdrl:Hsa.HRAS-mCherry)*^54^ in a Casper background ^55^ was used.

At 2.25 days post fertilization, zebrafish larvae were xenografted with MDA-MB-231 breast cancer cells. The MDA-MB-231 cells expressed *Ftractin-EGFP* stably to label the cells and reveal their actin organization^56^. The *Ftractin-EGFP* construct was a gift from Dr. Dyche Mullins at UCSF (Addgene plasmid # 58473) and MDA-MB-231 cells were a gift from Dr. Rolf Brekken (UT Southwestern Medical Center, TX). To study the effect of BLTP2 on migration and survival *in vivo*, we knocked down BLTP2 using shBLTP2 in MDA-MB-231 *Ftractin-EGFP* cells 5 days before xenografting as described above. An equal dose of control shUT virus was transduced into an equal number of MDA-MB-231 *F-tractin-EGFP* cells. We compared the outcome of the xenografts of shBLTP2 transduced cells with a xenograft of shUT transduced cells.

To xenograft the cells, they were grown up to 70-90% confluency and trypsinized for 3 min (Gibco, 15400-054). For injection, 4 × 10^6^ cells in 40 uL of cell culture media were prepared and stored on ice until xenografting into the yolk near the common cardinal vein of zebrafish larva. Injection was performed with glass capillary needles (World Precision Instruments, 1B100-4), pulled on a micropipette puller (Sutter Instrument, P-1000). Thereby, 50-500 cells were injected per fish. During the injection, zebrafish were anesthetized with Tricaine (Sigma Aldrich, E10521).

### In vivo cell migration and survival assay

To analyze the dissemination and survival of cancer cells *in situ*, we performed imaging of xenografted zebrafish. 5, 24, and 48 hours after injection, zebrafish xenografts were imaged on a Leica M205 FA fluorescence stereo microscope with a Planapo 1.0x objective and an X-Cite XYLIS illumination source. In addition, we obtained high-resolution images of cancer cells inside the vasculature in selected xenografts with axially swept light-sheet microscopy ^57^ on a custom microscope with an NA1.0 detection objective and 55x magnification. To immobilize the zebrafish for imaging, zebrafish embryos were anesthetized with 200 mg/l Tricaine (Sigma Aldrich, E10521) during imaging^58^. Between imaging sessions, zebrafish larvae were maintained at 34°C.

For each condition, 2 experimental repeats were performed with the shBTLP2 knockdown experiment containing n_1_=74 and n_2_=63 larvae; and shUT experiments contained n_1_=75 and n_2_=80 larvae.

To quantify cancer dissemination, we counted all zebrafish with micro-metastases to the tail part and with cancer cells remaining at the injection site in the yolk. We further counted all zebrafish with no cancer cells left and larvae that died during the experiment. To determine the spread of the cancer cells, we manually annotated the cancer cell locations of each zebrafish larva in a template fish using Fiji^59^. We obtained quantitative cancer dissemination maps by summing all manual annotations per time point and condition. To visualize the cancer dissemination maps, we overlaid them onto the template larva, with a rainbow look up table in Adobe Photoshop.

### Design of C. elegans strains, culture, and imaging

*C. elegans* strains used in this study: N2 Bristol(wildtype), AG674 *fmp-27Δ(av263) was* generated by CRISPR/Cas9 editing to delete full length of *bltp-2. C. elegans* strains were maintained with standard protocols in the Golden lab. The Bristol N2 strain was used as the wild type for CRISPR/Cas9 genome editing. 20 nucleotide sequence of the *fmp-27* specific crRNAs were selected with the help of a crRNA design tool from Integrated DNA Technologies (IDT). All crRNAs and tracrRNA were synthesized by Horizon Discovery (https://horizondiscovery.com). The single-stranded donor oligonucleotides (ssODNs) of the *bltp-2* repair template were synthesized by IDT, and the detailed sequence information of the CRISPR reagents is listed in Table 3. DIC images were taken by a spinning disk confocal system that includes a Photometrics Prime 95B EMCCD camera, and a Yokogawa CSU-X1 confocal scanner unit. Images were acquired by Nikon’s NIS imaging software using a Nikon 10x objective with 2-4 um z-step size; 10-20 planes were captured. The quantification of body length was analyzed by ImageJ/FIJI Bio-format plugin (National Institutes of Health) ^60, 61^. The animals were immobilized on 7% agar pads with an anesthetic (0.01% levamisole in M9 buffer). Statistical significance was determined by p-value from an unpaired 2-tailed t-test. Both the Shapiro-Wilk and Kolmogorov-Smirnov Normality tests indicated that all data follow normal distributions. We thank the Caenorhabditis Genetics Center, which is funded by the National Institutes of Health Office of Research Infrastructure Programs (P40OD010440), for providing strains for this study.

### Data presentation and statistics

Values were calculated and plotted in GraphPad Prism to generate histograms, violin plots, volcano plots, and heat maps. Experiments were conducted in at least triplicates. For statistical comparison, when two values were compared against each other, an unpaired t-test was conducted on GraphPad Prism to calculate and plot p-values in the graph. The p-values are indicated in the figures. To compare mean values of more than two entities, mean values were either compared against a common control or against each other using a one-way ANOVA and multiple comparison test on GraphPad Prism. The p-values are indicated in the figures.

